# STREME: Accurate and versatile sequence motif discovery

**DOI:** 10.1101/2020.11.23.394619

**Authors:** Timothy L. Bailey

## Abstract

Sequence motif discovery algorithms can identify novel sequence patterns that perform biological functions in DNA, RNA and protein sequences—for example, the binding site motifs of DNA- and RNA-binding proteins. The STREME algorithm presented here advances the state-of-the-art in *ab initio* motif discovery in terms of both accuracy and versatility. Using *in vivo* DNA (ChIP-seq) and RNA (CLIP-seq) data, and validating motifs with reference motifs derived from *in vitro* data, we show that STREME is more accurate, sensitive, thorough and rapid than several widely used algorithms (DREME, HOMER, MEME, Peak-motifs and Weeder). STREME’s capabilities include the ability to find motifs in datasets with hundreds of thousands of sequences, to find both short and long motifs (from 3 to 30 positions), to perform differential motif discovery in pairs of sequence datasets, and to find motifs in sequences over virtually any alphabet (DNA, RNA, protein and user-defined alphabets). Unlike most motif discovery algorithms, STREME accurately estimates and reports the statistical significance of each motif that it discovers. STREME is easy to use via its web server at http://meme-suite.org, and is fully integrated with the widely-used MEME Suite of sequence analysis tools, which can be freely downloaded at the same web site for non-commercial use.

## 1 Introduction

As do several existing motif discovery algorithms (e.g., Weeder [10], HOMER [5], STEME [12]), STREME makes use of a data structure called a generalized suffix tree [18]. It uses the generalized suffix tree to store the input sequences, and in the case of DNA, their reverse complements. Unlike Weeder and HOMER, however, which use the suffix tree to speed counting of approximate matches to individual words, STREME uses it to efficiently count matches to a position weight matrix (PWM) [15] representing a candidate motif. Another innovation of STREME is that, unlike HOMER and Weeder, which use separate searches of a suffix tree to find motifs with different widths, the STREME algorithm scores PWMs of all widths in the user-specified range in a *single* depth-first traversal of the suffix tree. This makes STREME faster, especially for wide motifs (width up to 30 positions), and more thorough in its search for the ideal width for each motif.

Like DREME [1] and HOMER, STREME evaluates motifs using a statistical test of the enrichment of matches to the motif in a primary set of sequences compared to a set of control sequences. STREME uses Fisher’s exact [3] test if the primary and control sequences have the same length distribution, and the Binomial test otherwise. Like DREME, and unlike HOMER, STREME can also discover motifs given just a primary set of sequences. In that case, STREME will create a control set by shuffling the letters of the primary sequences, preserving certain (user-specified) lower-order statistics of the sequences. Preserving the lower-order sequence statistics helps STREME avoid discovering uninteresting motifs. In addition, STREME always creates a Markov model of a user-specified order from the control sequences. STREME uses the Markov model in conjunction with the PWM when counting matches to the motif to further bias the search away from motifs that are mere artifacts of the lower-order statistics of the input sequences.

STREME (like DREME) reports accurate significance estimates for each motif that it discovers, in contrast to HOMER, Weeder and Peak-motifs [16], which do not report motif statistical significance, and MEME, whose motif significant estimates tend to be too conservative [9]. STREME accurately estimates motif statistical significance by evaluating the ability of each motif it discovers to classify primary and control sequences that it sets aside (holds out) and does not use during the motif discovery process.

STREME assumes that each primary sequence may contain zero or one occurrences (sites) of the motif (the so-called “ZOOPS” model [2]). Motif discovery will not be negatively affected if a primary sequence contains more than one occurrence of a motif, but unlike MEME, STREME cannot discover motifs given only a single primary sequence.

Finally, STREME will work with user-specified (“custom”) alphabets, as will MEME and DREME, but not the other algorithms studied here. This allows STREME to be applied to a much wider range of motif-discovery problems than HOMER, Peak-motifs or Weeder, including finding motifs in epigenetically modified DNA or post-translationally modified proteins,

In the remainder of this paper, we describe the STREME algorithm in more detail and present experimental results comparing its performance with several widely-used motif discovery algorithms. For the experimental comparisons, we consider motif discovery in ChIP-seq datasets (transcription factor (TF) binding motifs) and in CLIP-seq datasets (RNA-binding protein (RBP) motifs). In each case, we validate the predicted motifs using motifs derived using completely independent assays—high-throughput SELEX [6] (TF motifs) and RNAcompete [11] (RBP motifs). We also present an example of using STREME to find motifs in sequences in a user-defined alphabet (English).

## 2 Results

### 2.1 The STREME algorithm

STREME searches for motifs by performing step 1 (below), and then iterating steps 2 through 6 until the stopping criterion is met. The stopping criterion, chosen by the user, can be either a maximum number of motifs, or a minimum significance (maximum *p*-value) threshold.

#### 1. Dataset Preparation

STREME first reads the input sequence dataset(s) (primary, and optionally, control), converting to uppercase if the sequence alphabet is not case sensitive, and converting all ambiguous characters to a “separator” character that is not present in the alphabet.

To ensure that STREME will give the same results regardless of the order of the sequences in the input dataset(s), it sorts the input dataset(s) alphabetically by sequence content, and then randomizes the order of the sequences in the dataset(s).

Next, if the user does not provide a set of control sequences, STREME creates one from the primary sequences. Each primary sequence is shuffled, preserving the frequencies of all words of length *k* (“*k*-mers”) within it, where *k* can be specified by the user. The shuffling also preserves the positions of any separator characters. This prevents artifacts that can be caused by the presence of ambiguous characters in the sequences (such as the “N” character used by DNA repeat-masking programs).

STREME chooses the statistical test it will use. It will use Fisher’s exact test if the primary and control sequences have the same average length (within 0.01%), otherwise it will use the Binomial test.

Then, STREME creates a Markov model of the control sequences of order *k* – 1. STREME uses this model in conjunction with the PWM to compute the likelihood ratio scores of words during all stages of motif discovery and evaluation.

Finally, STREME creates a “hold-out” dataset for accurately assigning statistical significance to each discovered motif. By default, the hold-out set consists of a random sample of 10% of the sequences in the primary and control datasets.

#### 2. Suffix Tree Creation

STREME builds a generalized suffix tree that includes both the primary and control sequences (but not the hold-out set sequences). If the alphabet is complementable, STREME adds the reverse complement of each primary and control sequence to the tree as well. For the first round of STREME, the sequences are those created by Step 1, above. For subsequent rounds, the sequences from the last step (Step 6, below), with previous motifs erased, are used. STREME builds the suffix tree using code developed for MUMmer [7].

#### 3. Seed Word Evaluation

STREME uses the suffix tree to efficiently evaluate all words (up to a user-specified maximum length) that occur in the primary sequences, computing the *p*-value of each such word’s relative enrichment in the primary sequences using the chosen objective function. Each such word is called a “seed” word.

#### 4. Motif Refinement

STREME converts each of the four best seed words of each width in the user-specified range into a PWM (which we will refer to as the “motif”), and then iteratively refines each such motif. At each iteration of refinement, the current motif and the (*k* – 1)-order background are used with the suffix tree to efficiently find the best site in each sequence. The primary and control sequences are then sorted by the log-likelihood score of their best site, and the score threshold that optimizes the *p*-value of the statistical test is found. The iteration ends by using maximum likelihood estimation to estimate a new version of the motif from the single best site in each primary sequence whose score is above the optimal threshold. STREME uses this new motif in the next refinement iteration. Refinement stops when the *p*-value fails to improve, or the maximum allowed number of iterations (20) have been performed. As the final motif for the round, STREME selects the motif that best discriminates the primary sequences from the control sequences.

#### 5. Motif Significance Computation

STREME computes the statistical significance of the of the motif by using the motif and the optimal discriminative score threshold (based on the primary and control sequences) to classify the hold-out set sequences, and then applying the statistical test to the classification. Classification is based on the best match to the motif in each sequence (on either strand when the alphabet is complementable).

#### 6. Motif Erasing

STREME “erases” every site matching the best motif in the primary, control and hold-out sequences (and their reverse complements, if the alphabet is complementable) by converting the sites to the separator character. A site is considered a match if its log likelihood score according to the PWM exceeds the optimal score threshold determined in step 4. All matching sites (not just the best one in each sequence) are erased. To allow a certain amount of overlap between the sites of different motifs, STREME only sets letters in the site to the separator character if the letter’s likelihood ratio (according to the PWM) is positive.

### 2.2 Comparison of motif discovery algorithms on TF ChIP-seq data

#### 2.2.1 Accuracy

The accuracy of a motif discovery algorithm on a TF ChIP-seq dataset is its ability to discover an accurate version of the binding motif of the ChIP-ed TF in the set of DNA sequences identified as bound by the TF. We evaluate the accuracy of motif discovery algorithms by comparing their predicted motifs to the known motif for the transcription factor using the Tomtom motif comparison algorithm [4]. So that higher scores will correspond to more accurate motifs, we use minus the base-10 logarithm of the Tomtom *p*-value of the similarity between the discovered motif and the reference motif as our motif accuracy score. To avoid circularity, we use motifs derived using SELEX—an *in vitro* assay—as our reference motifs. For each of 40 ENCODE TF ChIP-seq experiments in K562 cells for which there is a known TF motif in the Jolma *et al*. [6] compendium of SELEX-derived motifs, we prepare primary and control datasets, and assign a single SELEX motif to be the reference motif for that experiment (see Methods, below). The primary sequence dataset for an experiment consists of the 100bp sequence regions centered on each of the ChIP-seq peaks; the control dataset comprises a shuffled version of each of the primary sequences, with the frequency of 3-mers in each sequence preserved. (Weeder does not use a control dataset, so we present it with only the primary dataset.) Because ChIP-seq peak regions often contain enriched motifs besides that of the ChIP-ed TF (e.g., binding motifs of cofactors), the motif discovery algorithms were each allowed to report five motifs when presented with the primary and control datasets. We run each algorithm with its default settings, except we set the minimum and maximum motif widths to 8 and 12, respectively. For the three algorithms that construct a background model from their input sequences (STREME, MEME and Peakmotifs), we specify that a second order Markov model be used.

As shown in Fig. 1a, the accuracy of STREME is as good or better than that of the other algorithms tested here. The best motif found by STREME in ChIP-seq data is more similar to the reference motif derived from SELEX data, on average, than the best motif found by the other algorithms, across a wide range of accuracy score thresholds. STREME, MEME and HOMER all discover a motif with an accuracy score of at least 5 (Tomtom *p*-value ≤ 10^−5^) in at least 70% of the ChIP-seq datasets. STREME actually finds such a motif 82.5% of the time. Additionally, STREME discovers about twice as many highly accurate motifs as the other two algorithms at the more stringent motif accuracy score threshold of 9 (Tomtom *p*-value ≤ 10^−9^). At this motif accuracy score threshold, STREME is successful on 32.5% of the ChIP-seq datasets, whereas HOMER is only successful on 12.5% of the datasets.

**Figure 1:**
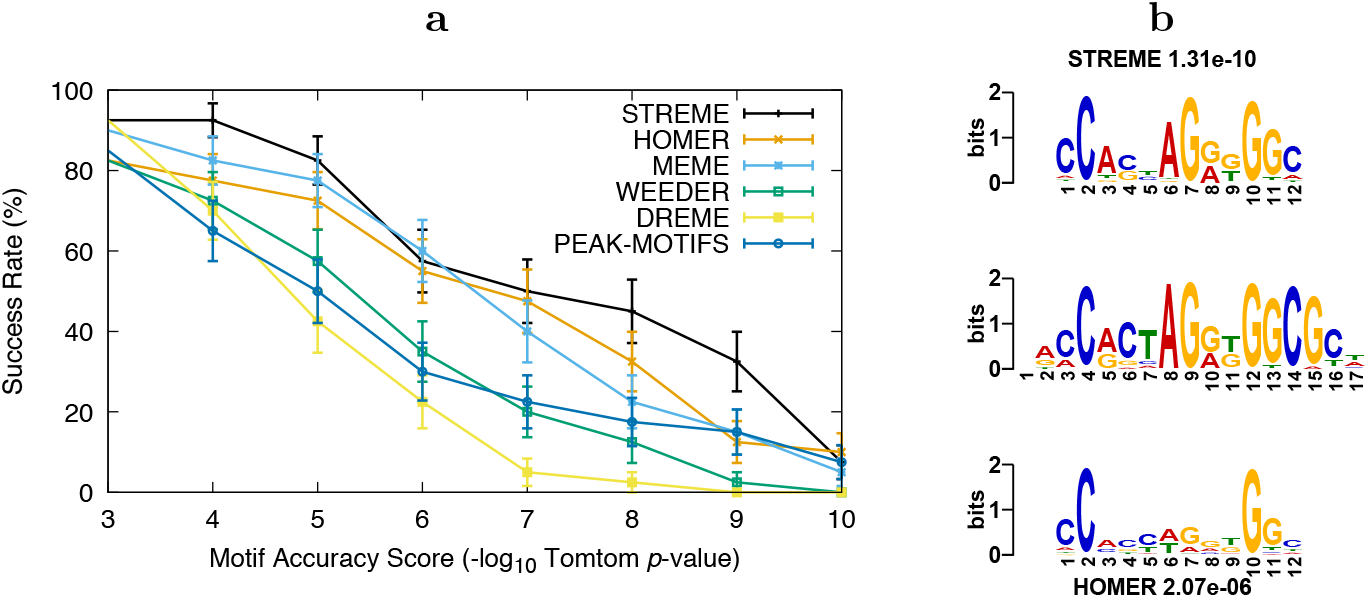
Accuracy of motif discovery algorithms on ENCODE TF ChIP-seq datasets. The curves in Panel **a** show the percentage of times *(Y*) the best motif found by the named algorithm has motif accuracy score ≥ *X*, averaged over 40 ChIP-seq datasets. Panel **b** shows the sequence logos and accuracies of the best motifs (and their accuracies) found by STREME (top) and HOMER (bottom) in an ENCODE ChIP-seq dataset (UtaK562Ctcf), aligned to the SELEX motif CTCF_full (center) from Jolma *et al*. [6] that we used to evaluate those motifs.

Fig. 1b gives an example of a type of motif where the HOMER algorithm is sometimes less accurate than STREME because, unlike STREME, HOMER does not keep track of strand information for individual motif sites. This causes HOMER to sometimes combine overlapping sites on the two DNA strands into a motif that is a perfect palindrome, when, in fact the motif is quite asymmetrical (e.g., Fig. 1b). STREME avoids this problem by keeping track of strand information, and by never allowing more than one site in a sequence (on either strand) to contribute to a motif. Sequence logos for the best STREME and HOMER motifs are shown, along with their motif accuracies (Tomtom *p*-values), for all 40 TF ChIP-seq datasets studied here in Table 1. As seen in that table, the STREME motif is more accurate than the HOMER motif for 29 of the 40 TF ChIP-seq datasets.

**Table 1:**
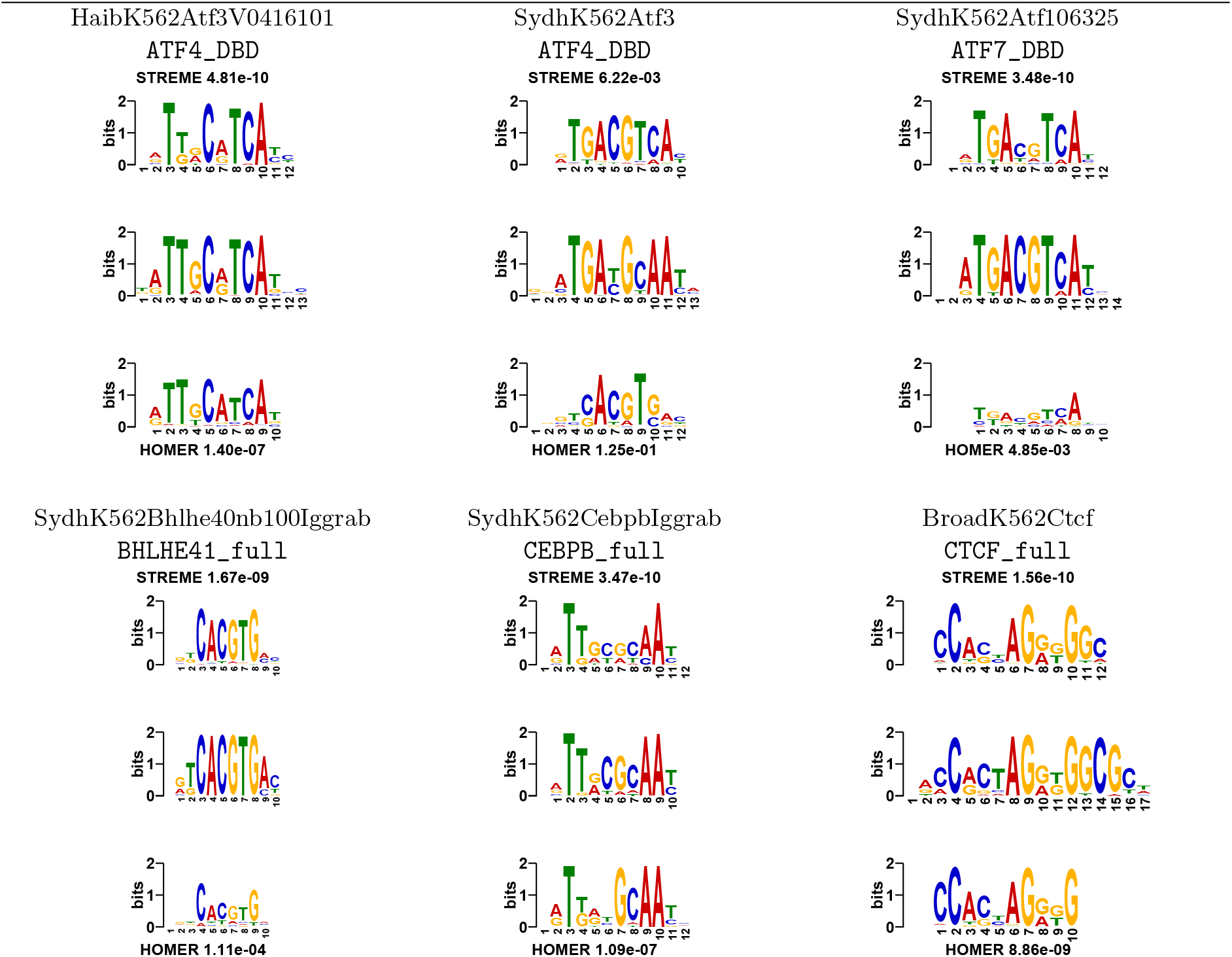
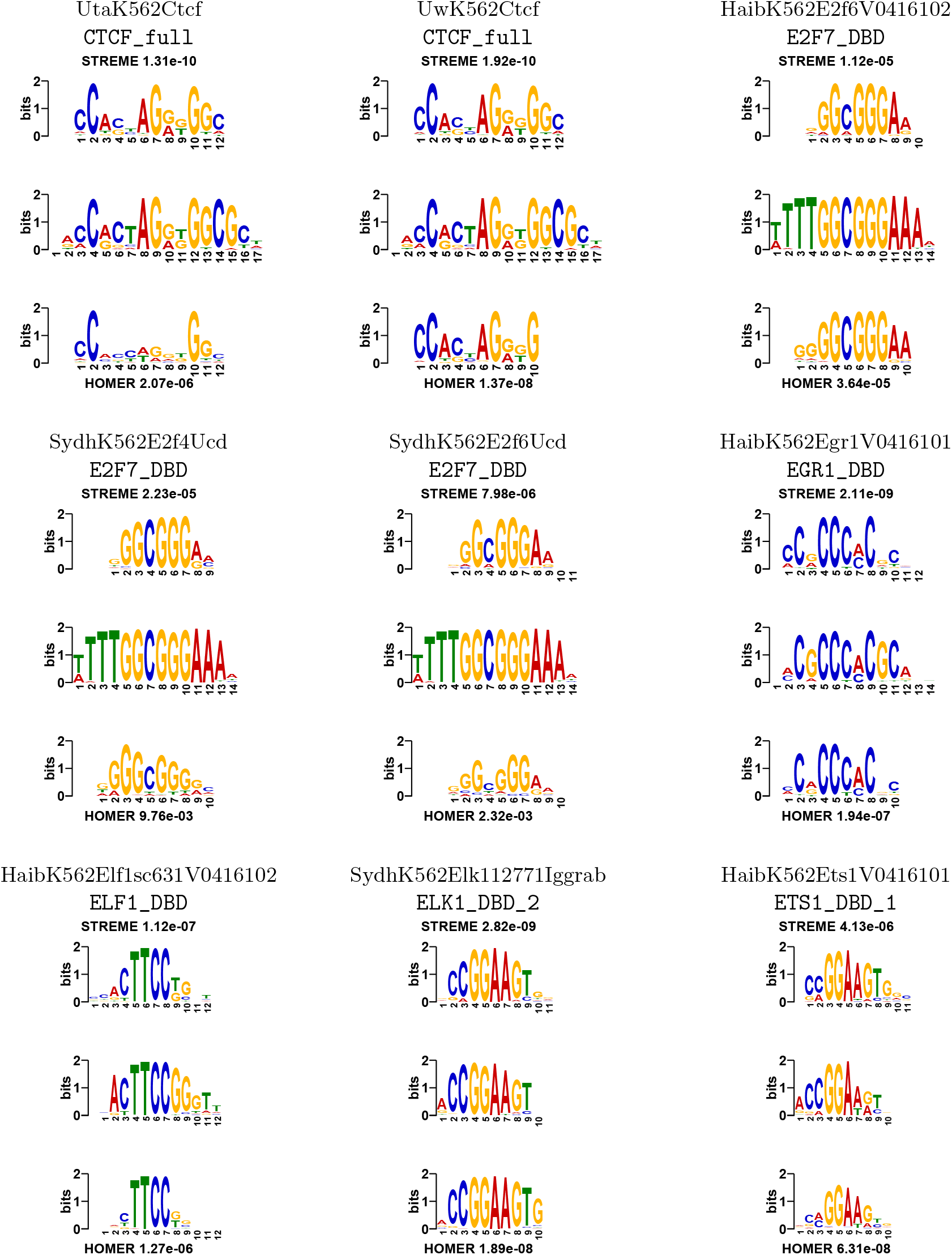
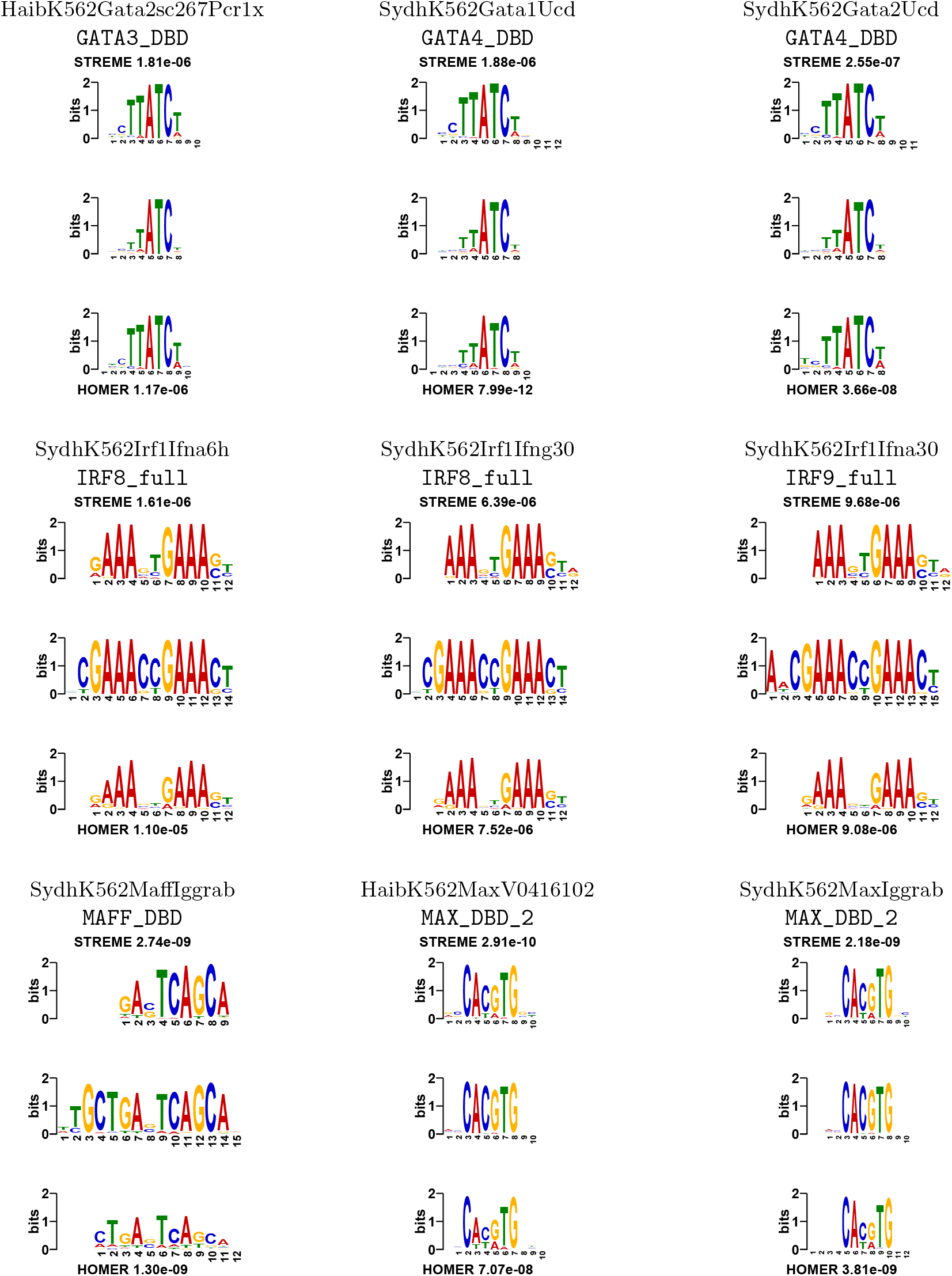
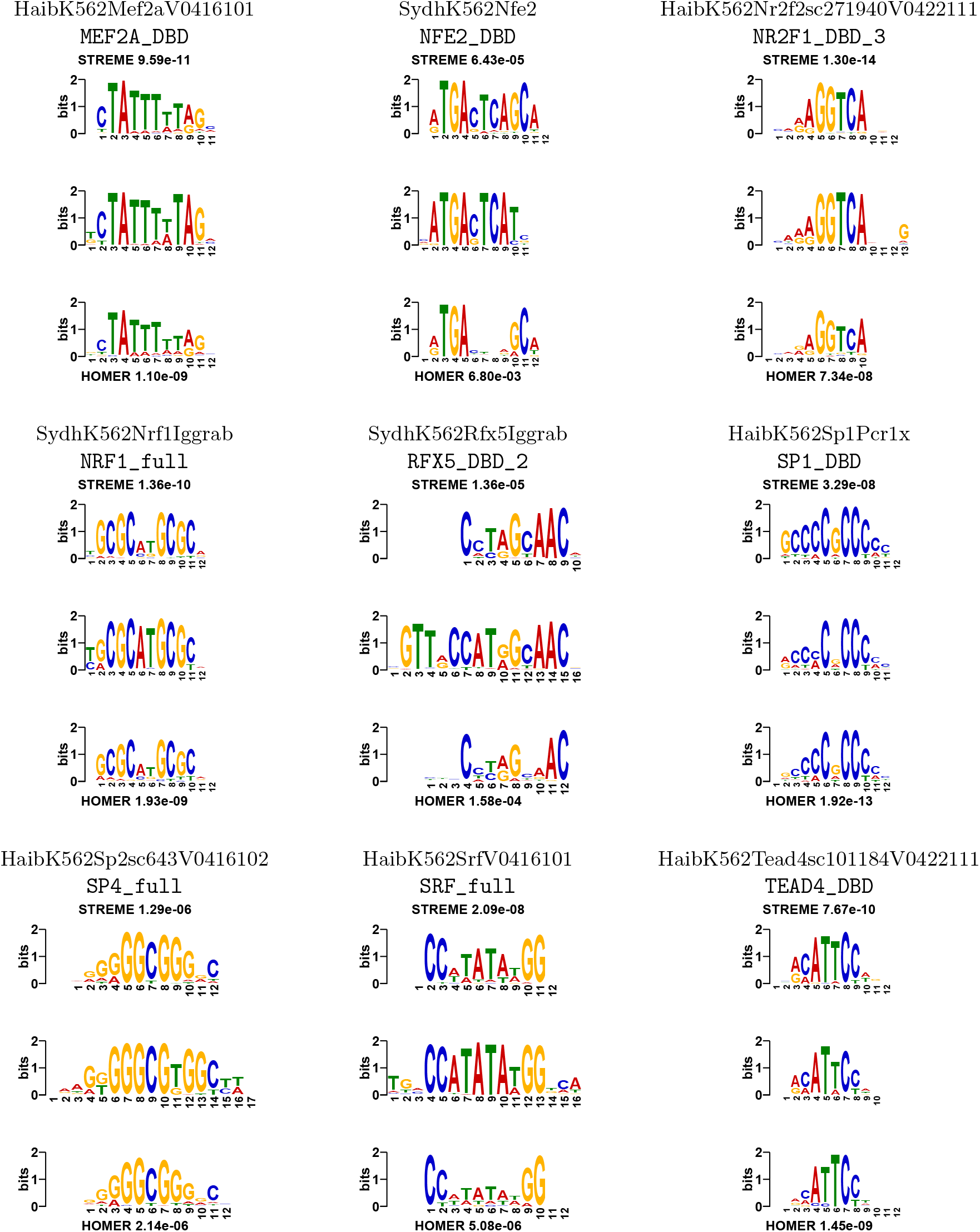
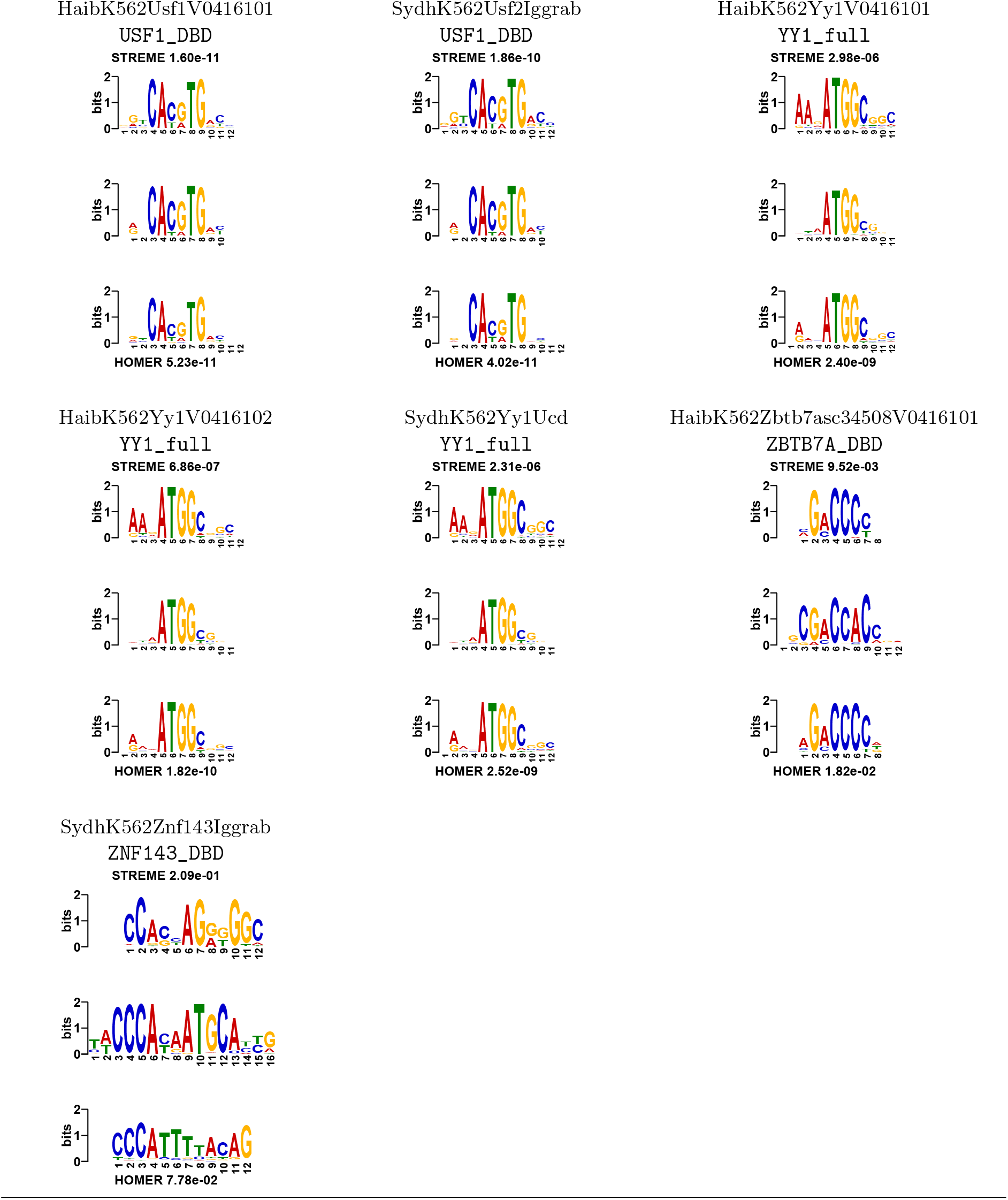
Comparison of STREME and HOMER ChIP-seq motifs. The top two lines of each panel show the name of the ENCODE TF ChIP-seq dataset and the name of the SELEX motif (from Jolma *et al*. [6]) that we used for evaluating that experiment. The three sequence logos show the STREME motif (top) and the HOMER motif (bottom) aligned to the reference motif (center). Above and below the aligned logos are the names of the two algorithms and the accuracy (Tomtom *p*-value) of the motif discovered by the named algorithm. (A smaller Tomtom *p*-value indicates that the discovered motif is more similar to the reference motif.) The motif found by STREME is more accurate than that found by HOMER for 29 out of the 40 TF ENCODE ChIP-seq datasets (done in K562 cells) for which there is a high-throughput SELEX reference motif.

#### 2.2.2 Sensitivity

The sensitivity of a motif discovery algorithm is its ability to discover motifs that are present in only a small fraction of the primary input sequences. To study this aspect of performance, we construct a series of progressively more difficult primary datasets from each of the original 40 primary TF ChIP-seq datasets by shuffling the letters of up to 99% of the sequences in the dataset, preserving the frequency of 3-mers, which removes any motif occurrences from the shuffled sequences. As before, for each such “diluted” primary dataset, we create a control dataset by shuffling the letters of all the sequences in it, preserving the frequency of 3-mers. We let each motif discovery algorithm report five motifs, allowing the algorithm to choose the optimal motif width in the range 8 to 12.

To analyze the results, we must choose a motif accuracy score threshold for deciding if the algorithm has found a motif matching the reference motif for the dataset. Fig. 2 compares the algorithms using an motif accuracy score threshold of 5 (Tomtom *p*-value ≤ 10^−5^); using higher or lower thresholds gives similar results (data not shown). Across the range of dataset “purity” from 100% (the original ChIP-seq peak sequences) to 1% (99% of sequences are shuffled), STREME is at least as likely to discover the ChIP-ed TF’s motif as HOMER, and better than the other algorithms. We conclude that the sensitivity of STREME is as good or better than that of the algorithms we examine here.

**Figure 2:**
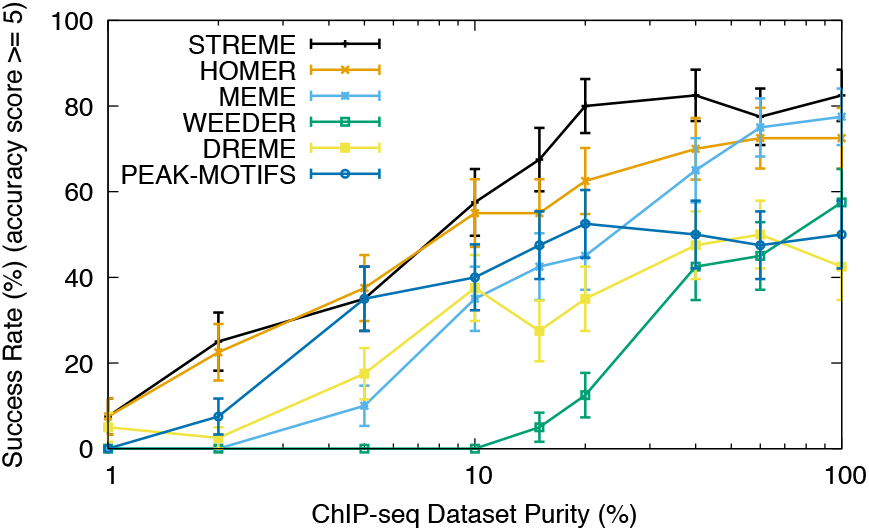
Sensitivity of motif discovery algorithms on ENCODE TF ChIP-seq datasets. Each point shows the percentage of times (*Y*) the best motif found by the named algorithm in a primary dataset that has been diluted to a given purity (*X*) has motif accuracy score at least 5 (Tomtom *p*-value ≤ 10^−5^), averaged over 40 ChIP-seq datasets.

#### 2.2.3 Thoroughness

The thoroughness of a motif discovery algorithm is the degree to which it can discover many distinct motifs in a given primary set of sequences. This is a particularly important aspect of motif discovery in TF ChIP-seq datasets, where multiple TFs often bind in close proximity to regulate transcriptional expression. To measure the ability of STREME and other algorithms to discover multiple motifs, we construct an artificial ChIP-seq dataset by combining 100 randomly-selected sequences from each of 21 of our 40 ENCODE TF ChIP-seq primary datasets. (See Methods, below, for how the 21 datasets were selected.) This creates a set of primary sequences where fewer than 5% contain peak regions for a given ChIP-ed TF, and where there are sequences containing binding sites for all 21 ChIP-ed TFs. We repeat the random sampling process 20 times to create 20 distinct primary datasets. Using these 20 primary datasets and shuffled versions of them as control datasets, we allow each motif discovery algorithm to report 25 motifs with widths in the range 8 to 12, and we measure each algorithm’s average success rate at finding all 21 reference motifs at different motif accuracy score thresholds.

Fig. 3 shows that STREME is generally more thorough in this setting than the other algorithms tested here. At motif accuracies less stringent than *p* = 10^−7^ (accuracy score < 7), STREME finds substantially more motifs than any of the other algorithms. At stricter accuracy thresholds, only MEME is as thorough as STREME.

**Figure 3:**
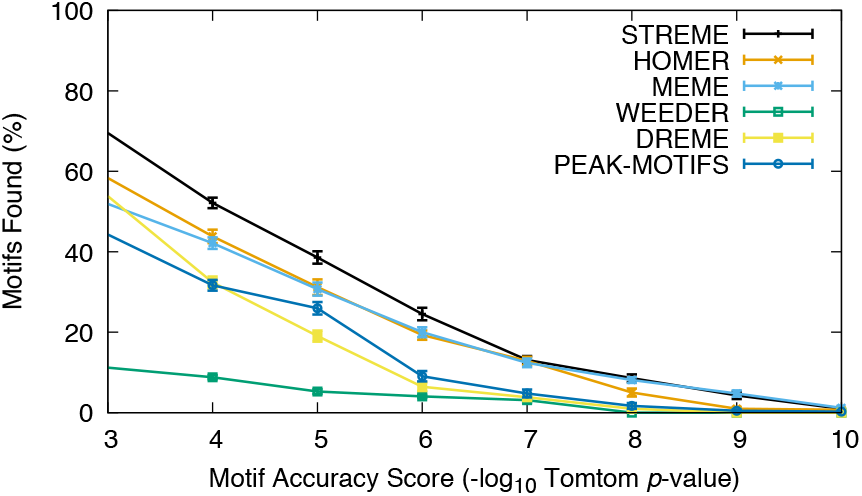
Thoroughness of motif discovery algorithms on combined ENCODE TF ChIP-seq datasets. The curves show the percentage of 21 reference motifs (*Y*) for which the named algorithm finds a motif matching it with given motif accuracy score (*X*) or better, averaged over 20 combined datasets.

#### 2.2.4 Speed

Fig. 4 shows the running time of the motif discovery algorithms on each of the 40 experiments described above in the Accuracy section. We run the algorithms using a single thread on a 4.0 GHz Intel Core i7 processor with 16GB of memory. Only Peak-motifs is faster than STREME, and only when the primary sequence dataset contains more than 20,000 sequences of 100bp. With fewer than 10,000 sequences, STREME is up to an order of magnitude faster than Peak-motifs. STREME is slightly faster in this setting than HOMER, and much faster than the other algorithms tested here. As we shall show below (see Fig. 6), STREME is much faster than HOMER for finding motifs wider than 12 positions. At all dataset sizes, STREME is approximately an order of magnitude faster than DREME, the algorithm it is designed to replace within the MEME Suite.

**Figure 4:**
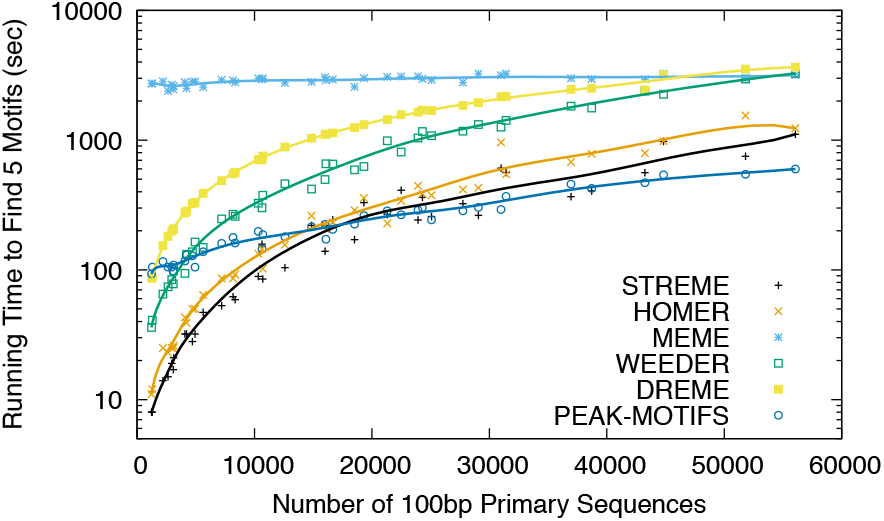
Speed of motif discovery algorithms on ENCODE TF ChIP-seq datasets. Each point represents the running time (*Y*) of the named algorithm on one of 40 ENCODE TF ChIP-seq datasets containing the given number of sequences (*X*). For ease of interpretation, the points for each algorithm have been fit with a smooth Bezier curve.

#### 2.2.5 Choosing the background model

In the preceding sections we use a value of *k* = 3 when creating sets of control sequences from each primary set by shuffling each sequence while preserving the frequencies of all *k*-mers within it. For STREME, MEME and Peak-motifs, we also instruct them to build an internal Markov model of order *k* – 1 = 2.

This choice of *k* is justified by the results shown in Fig. 11, which shows how each of the algorithms studied here (except Weeder) performs in terms of accuracy, sensitivity and thoroughness using values of *k* from 1 to 4. (Weeder is not included in the figure as it does not use a set of control sequences.) For all algorithms tested here, using a value of *k* = 3 provides (near) optimum results. Fig. 11 shows that, with the TF ChIP-seq datasets we use here, the effect of *k* is particularly large for STREME and HOMER.

#### 2.2.6 Accuracy on “small” datasets

We also study the accuracy of motif discovery algorithms on datasets with from 10 to 1000 sequences, using randomly chosen samples of increasing size from each of the 40 ENCODE TF ChIP-seq datasets. From each of these primary sequence datasets, we create a control dataset by randomly shuffling each sequence while preserving the frequencies of 3-mers. We run the motif discovery algorithms on each of the resulting pairs of datasets, and measure the accuracy of the best motif out of five, as before (see the subsection on “Accuracy”, above).

Fig. 5 shows, that with small datasets, the MEME algorithm, which is based on maximizing motif information content, more frequently finds a motif with accuracy score at least 5 (Tomtom *p*-value ≤ 10^−5^) than the other algorithms, all of which are based on maximizing the motif’s ability to classify sequences. In this setting, STREME performs the best of the classification-based algorithms, substantially better than HOMER. These observations remain true using motif accuracy score thresholds from 4 to 7, inclusive (data not shown).

**Figure 5:**
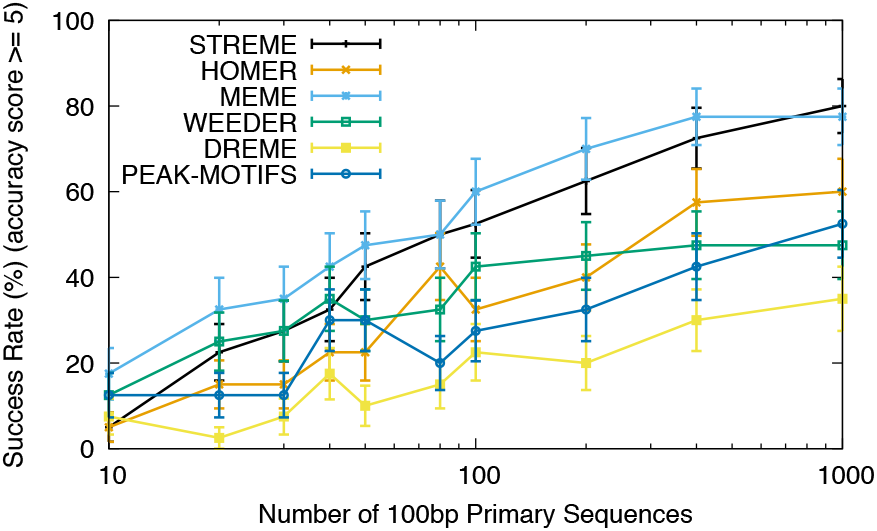
Accuracy of motif discovery algorithms on “small” datasets. Each point shows the percentage of times (*Y*) the best motif found by the named algorithm in a (sampled) dataset with the given number of sequences (*X*) has motif accuracy score at least 5 (Tomtom *p*-value ≤ 10^−5^), averaged over downsamples of each of 40 TF ChIP-seq datasets.

With primary datasets containing more than 1000 sequences, MEME bases its search on only a random subsample of 1000 sequences. (It does this because the running time of MEME’s underlying algorithm increases at least quadratically with the size of the sequence dataset.) With full-size ChIP-seq datasets, which typically contain many thousands of sequences, the classification-based algorithms (especially STREME) find more accurate motifs (Fig. 1), find fainter motifs (Fig. 2), and find more co-factor motifs (Fig. 3).

#### 2.2.7 Performance finding “wide” motifs

Many TF binding motifs are wider than 12 positions, the maximum motif width we set for the motif discovery algorithms in the results presented thus far. For example, the SELEX reference motifs for ATF4, ATF7, E2F7, EGR1, IRF8, MAFF, N2F1, RFX5, SP4, and ZNF143 all have widths between 13 and 17 (see Table 1). Three of the motif discovery algorithms studied here (STREME, HOMER and MEME) allow the user to specify the range of motif widths that the algorithm may search for. We conduct additional experiments to determine how running time scales with maximum motif width (up to a maximum of 30), and how the accuracy, sensitivity and thoroughness of the algorithms on TF ChIP-seq datasets is affected when we set the maximum motif width to 18 (instead of 12).

Fig. 6 shows how the running times of the three motif discovery algorithms scale with maximum motif width using randomly chosen DNA sequences. We run each of the algorithms on 25 (artificial) primary sequence datasets, with the maximum motif width specified as a number from 10 to 30, and the minimum motif width always set to 8. We create each primary sequence dataset by sampling one of our 40 ENCODE TF ChIP-seq datasets. Each such artificial sequence dataset contains 10,000 length 100bp DNA sequences, and we create a matching control dataset by shuffling the primary dataset while preserving the frequencies of 3-mers.

**Figure 6:**
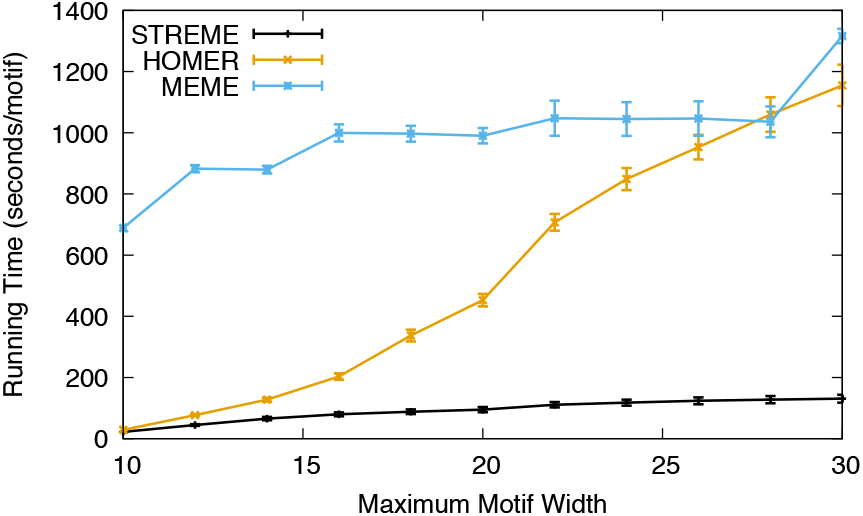
Running time as a function of maximum motif width. Each point shows the running time in seconds per motif found (*Y*) when the named motif finder is run with the given maximum motif width (*X*) set, averaged over 25 ChIP-seq primary sequence datasets each containing 10,000 sequences of length 100bp. Error bars show standard error. The points for a given motif discovery algorithm are connected with straight lines for ease of interpretation. The algorithms were run on a 3.2 GHz Intel Core i7 processor with 16GB of memory.

STREME is much faster than either HOMER or MEME for finding motifs wider than 12, as seen in Fig. 6. For finding motifs up to 30 positions wide, STREME is almost an order of magnitude faster than the other two algorithms. The improved speed relative to HOMER is due to the way STREME searches its suffix tree, computing scores for motifs from the minimum to the maximum specified width in a single, depth-first traversal of the tree. By contrast, HOMER repeats its traversal of the tree for each width in the width range. The running time of MEME with large datasets such as used here (10,000 sequences) is generally much higher than that of HOMER, except for the widest motifs.

Next we study how the accuracy, sensitivity, thoroughness and speed of the motif discovery algorithms compare when they search for wider motifs in the same 40 ENCODE TF ChIP-seq datasets we used in the previous sections. The only change to the experiments from those sections is that we ran STREME, HOMER and MEME with the maximum motif width parameter of each set to 18, rather than 12.

Fig. 7a shows that STREME finds more motifs across all motif accuracy score thresholds than either HOMER or MEME even with the wider maximum motif setting. Similarly, Fig. 7b shows that STREME is still at least as sensitive as HOMER, and considerably more so than MEME, even with the maximum motif width set to 18 instead of 12. STREME is also still more thorough than HOMER and MEME when we set the motif accuracy score threshold below 7 (Fig. 7c). When searching for motifs with widths up to 18, STREME is about an order of magnitude faster than HOMER (Fig. 7d).

**Figure 7:**
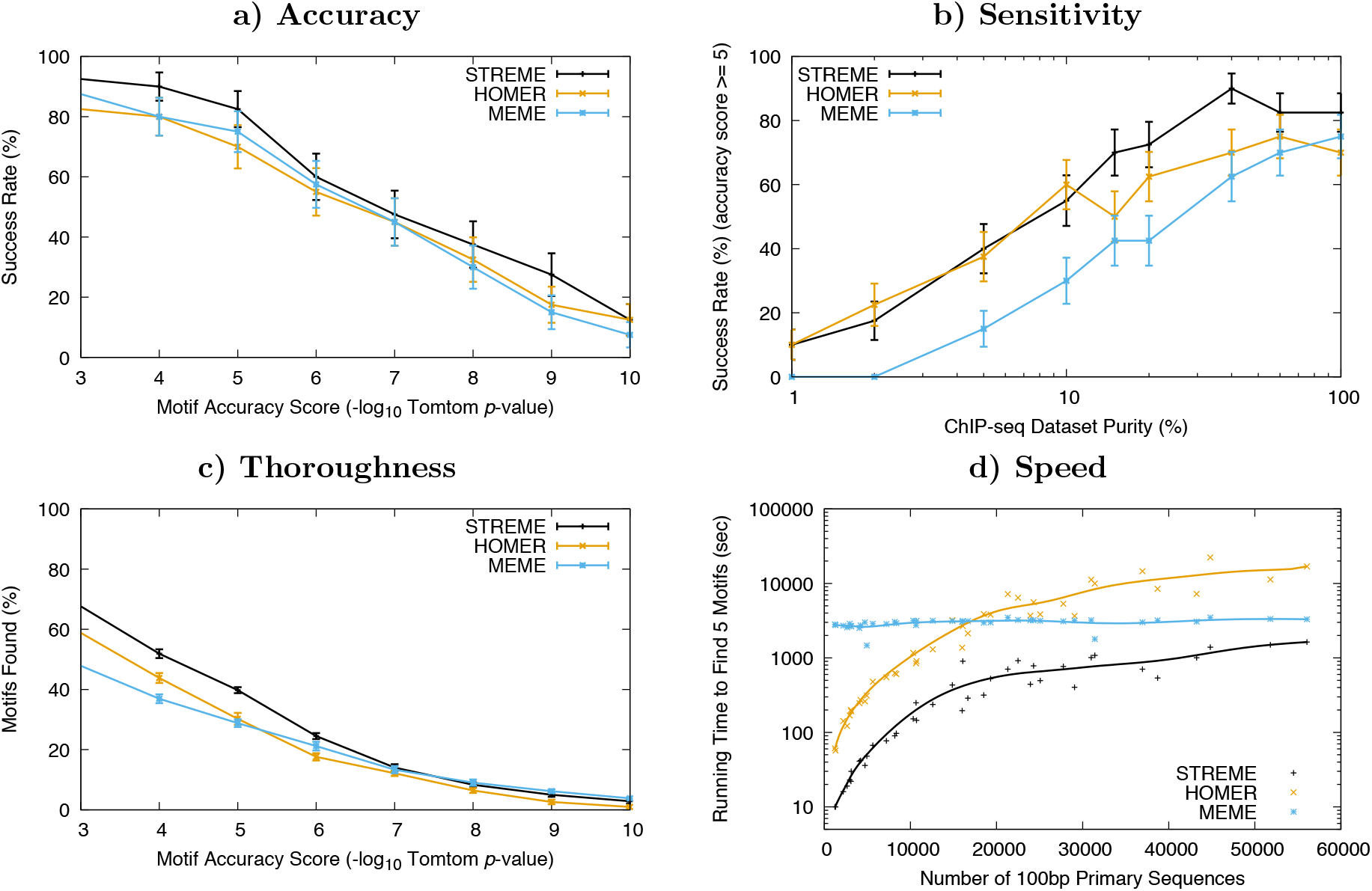
Performance discovering wide motifs (8 ≤ width ≤ 18) in TF ChIP-seq datasets. The curves in Panel **a** show the percentage of times (*Y*) the best motif found by the named algorithm has accuracy *X* or better, averaged over 40 TF ChIP-seq datasets. Each point in Panel **b** shows the percentage of times (*Y*) the best motif found by the named algorithm in a primary dataset that has been diluted to a purity *X* has accuracy *p* < 10^−5^, averaged over 40 TF ChIP-seq datasets. The curves in Panel **c** show the percentage of 21 reference motifs (*Y*) for which the named algorithm finds a motif matching it with motif accuracy score *X* or better, averaged over 20 combined datasets, each constructed from 100 randomly-selected sequences from each of 21 TF ChIP-seq datasets. Each point in Panel **d** represents the running time (*Y*) of the named algorithm on one of 40 ENCODE TF ChIP-seq datasets containing *X* sequences. Running times are on a 4.0 GHz Intel Core i7 processor with 16GB of memory, and, for ease of interpretation, the points for each algorithm have been fit with a smooth Bezier curve.

### 2.3 Comparison of motif discovery algorithms on CLIP-seq datasets

The accuracy of a motif discovery algorithm on an RNA-binding protein (RBP) CLIP-seq dataset is its ability to discover an accurate version of the binding motif of the RBP in the set of RNA sequences identified as bound by the RBP. As with the DNA (ChIP-seq) experiments above, we compare motif discovery algorithms run on data from an *in vivo* assay—eCLIP data from ENCODE [17]—using reference motifs for the same RBPs derived using an *in vitro* assay— RNAcompete [11]. As before, we use the Tomtom motif comparison algorithm to estimate the accuracy of the discovered motifs. We identify twenty eCLIP datasets for which an RNAcompete motif exists for the same RBP. As the primary sequence dataset, we use the full-length RNA sequences identified as bound by the ENCODE eCLIP experiment. We find that using shuffled sequences as the control dataset gives poorer results (data not shown) than using a random sample of 10,000 full-length bound sequences identified in the 120 ENCODE RNAcompete experiments, and that is what we use in the following experiments. We let each motif discovery algorithm discover five motifs, and allow the width to range from 7 to 8, since that is the range of widths in the RNAcompete RBP motif compendium. As with the DNA (ChIP-seq) experiments above, for the three algorithms that construct a background model from their input sequences (STREME, MEME and Peak-motifs), we instruct them to build a second order Markov model.

Fig. 8 shows that none of the motif discovery algorithms finds a motif that matches the reference motif more than 50% of the time even at a very permissive motif accuracy score threshold of 3 (Tomtom *p*-value ≤ 0.001). STREME and Weeder are perhaps slightly more accurate than the others, but the sample size is too small to declare that the difference is significant. Table 2 shows sequence logos [13] for the the best STREME and Weeder motifs, and their accuracies (Tomtom *p*-values), aligned to the logo of the reference motif, for each of the 20 eCLIP experiments.

**Figure 8:**
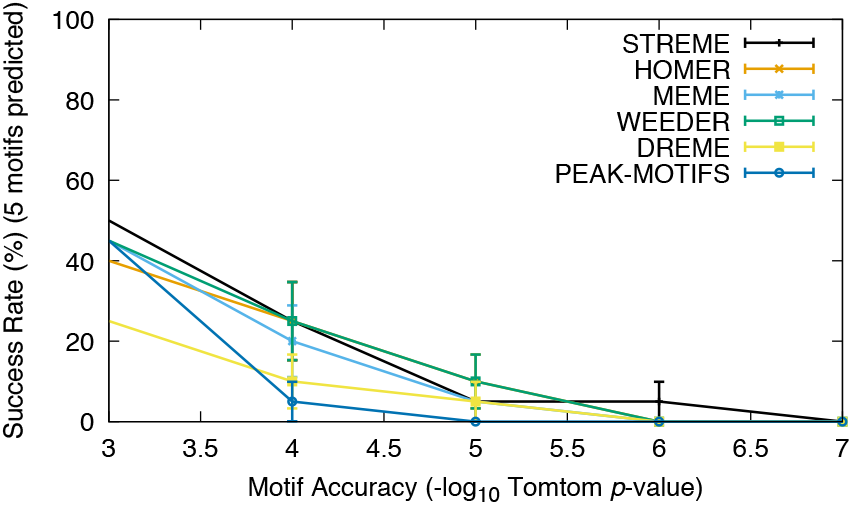
Accuracy of motif discovery algorithms on ENCODE RNA-binding protein eCLIP datasets. The curves in Panel **a** show the percentage of times (*Y*) the best motif found by the named algorithm has motif accuracy score *X* or better, averaged over 20 eCLIP datasets.

**Table 2:**
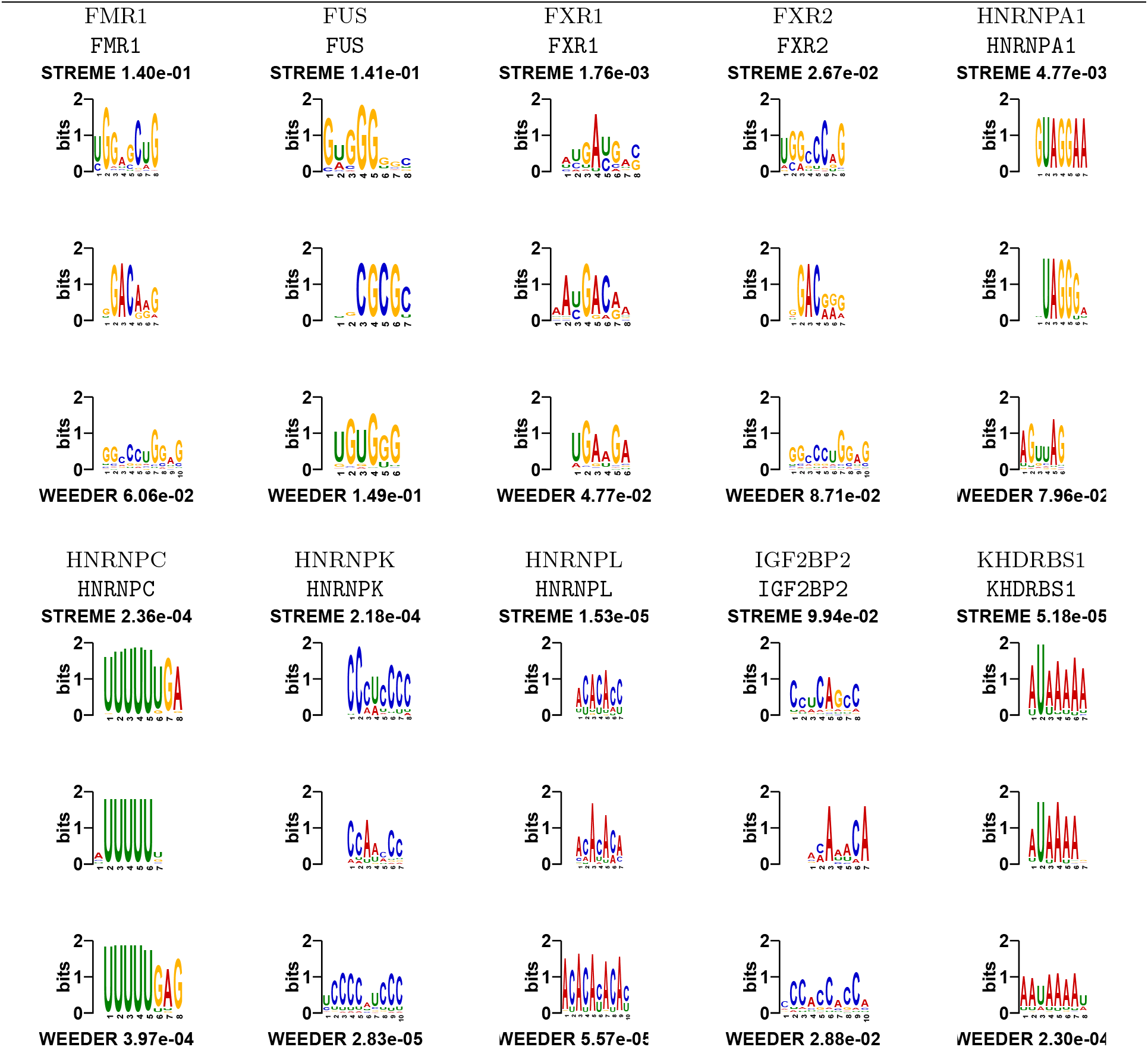
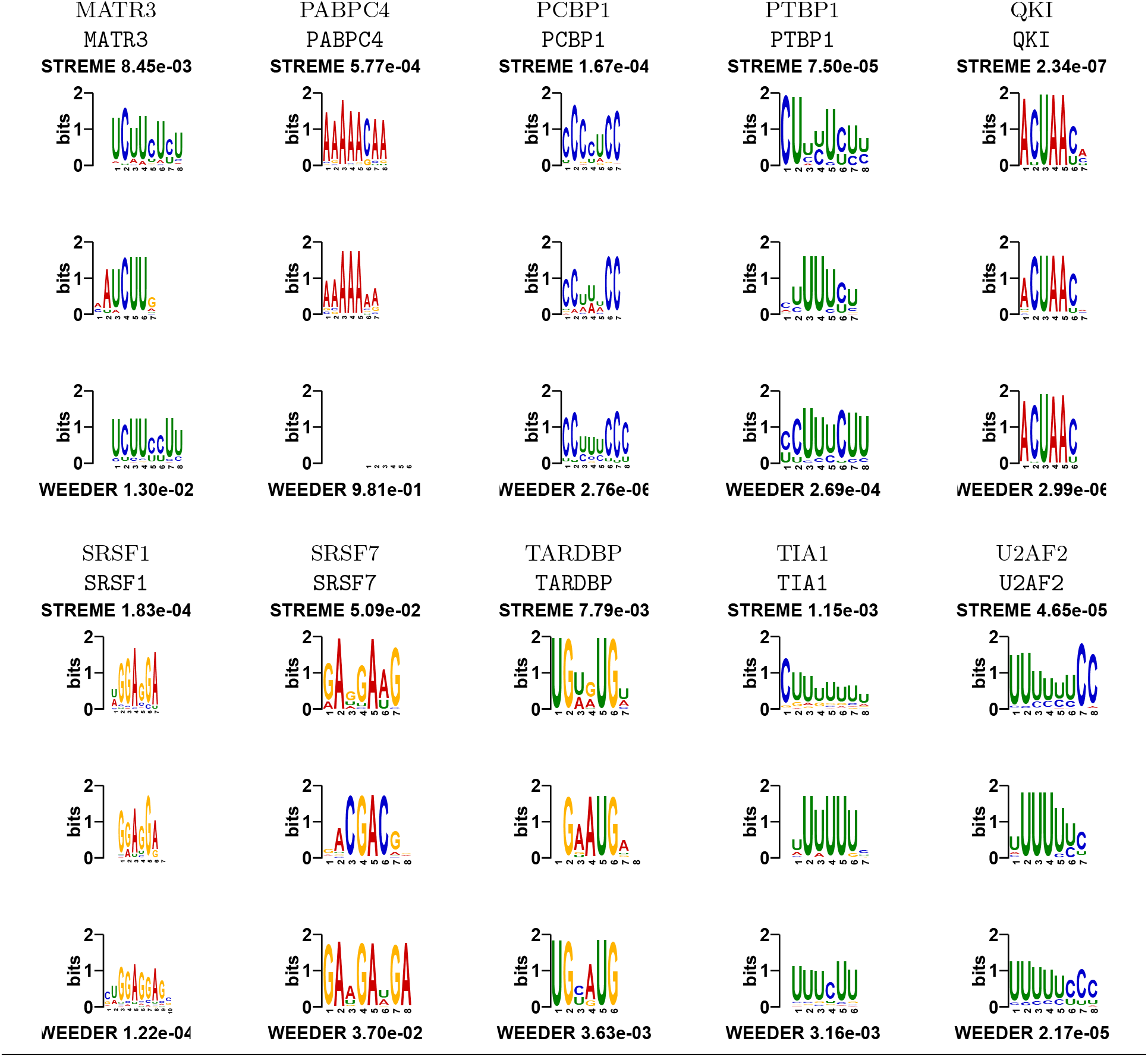
Comparison of STREME and Weeder eCLIP motifs. The top two lines of each panel show the name of the ENCODE RNA-binding protein eCLIP dataset and the name of the RNAcompete motif (from Ray *et al*. [11]) that we used for evaluating that experiment. The three sequence logos show the STREME motif (top) and the Weeder motif (bottom) aligned to the reference motif (center). Above and below the aligned logos are the names of the two algorithms and the accuracy (Tomtom *p*-value) of the motif discovered by the named algorithm. (A smaller Tomtom *p*-value indicates that the discovered motif is more similar to the reference motif.)

The lower motif accuracy we observe in this experiment is partly due to the fact that the RBP reference motifs are generally much shorter than TF binding motifs, which causes the Tomtom *p*-values of even very accurate motifs to be higher. In addition, in the three (out of 20) datasets where both STREME and Weeder fail to find a motif that is at all similar to the reference motif (FMR1, FUS, IGF2B2 in Table 2), the problem might lie with the algorithms, with those particular eCLIP datasets, or with the RNAcompete reference motifs. To check the first possibility, we compared the statistical enrichment of the RNAcompete reference motif and the most similar STREME motif for each experiment using the ENR motif enrichment algorithm from the MEME Suite. The motifs found by STREME and Weeder in the FMR1, FUS and IGF2BP2 eCLIP datasets are much more enriched than the reference motifs in those datasets (data not shown). For only two of the 20 eCLIP experiments was the RNAcompete reference motif more enriched in the eCLIP sequences than the STREME motif. The same was true for the motifs found by Weeder—in only two out of 20 cases was the RNAcompete motif more enriched than the Weeder motif. These results suggest that there may be problems with either the *in vitro* RNAcompete reference motifs or with the *in vivo* eCLIP datasets for FMR1, FUS and IGF2BP2.

### 2.4 Finding motifs using custom alphabets—a “novel” application

To illustrate the versatility of STREME due to its ability to use user-defined alphabets, we used it to discover the most distinguishing words in Moby Dick, a novel by Herman Melville. In many ways, this task is similar to discovering biologically important sequence motifs in biological sequences. To make the task more similar to biological motif discovery, and to make it more difficult, we remove all capitalization, spaces and punctuation from the text of the novel, and we randomly “mutate” 20% of the letters in the text to a (randomly-chosen) different letter from the English alphabet. We then create a FASTA file from the processed text, where each sentence in the novel becomes a FASTA sequence. The result is a FASTA file with 18,371 sequences ranging in length from 5 to 64 (mean length 51), containing 936,096 sequence characters in total. We also create an alphabet definition file in the simple format required by STREME, which consists of the line “`ALPHABET English`” followed by each of the 26 English letters on its own line. Although not needed here, the alphabet definition file can also specify a name and a color for each letter, as well as pairs of letters that are considered complements (as for DNA bases).

When we run STREME on the resulting FASTA file to search for motifs of widths from 3 to 12 using shuffled control sequences preserving words of width 2 (2-mers), it discovers 114 motifs with *p*-values less than 0.05. STREME always continues searching until the last three motifs found do not satisfy the user-specified significance level, and Table 3 shows the names, *p*-values and sequence logos of all 117 motifs that STREME finds in this dataset.

**Table 3:**
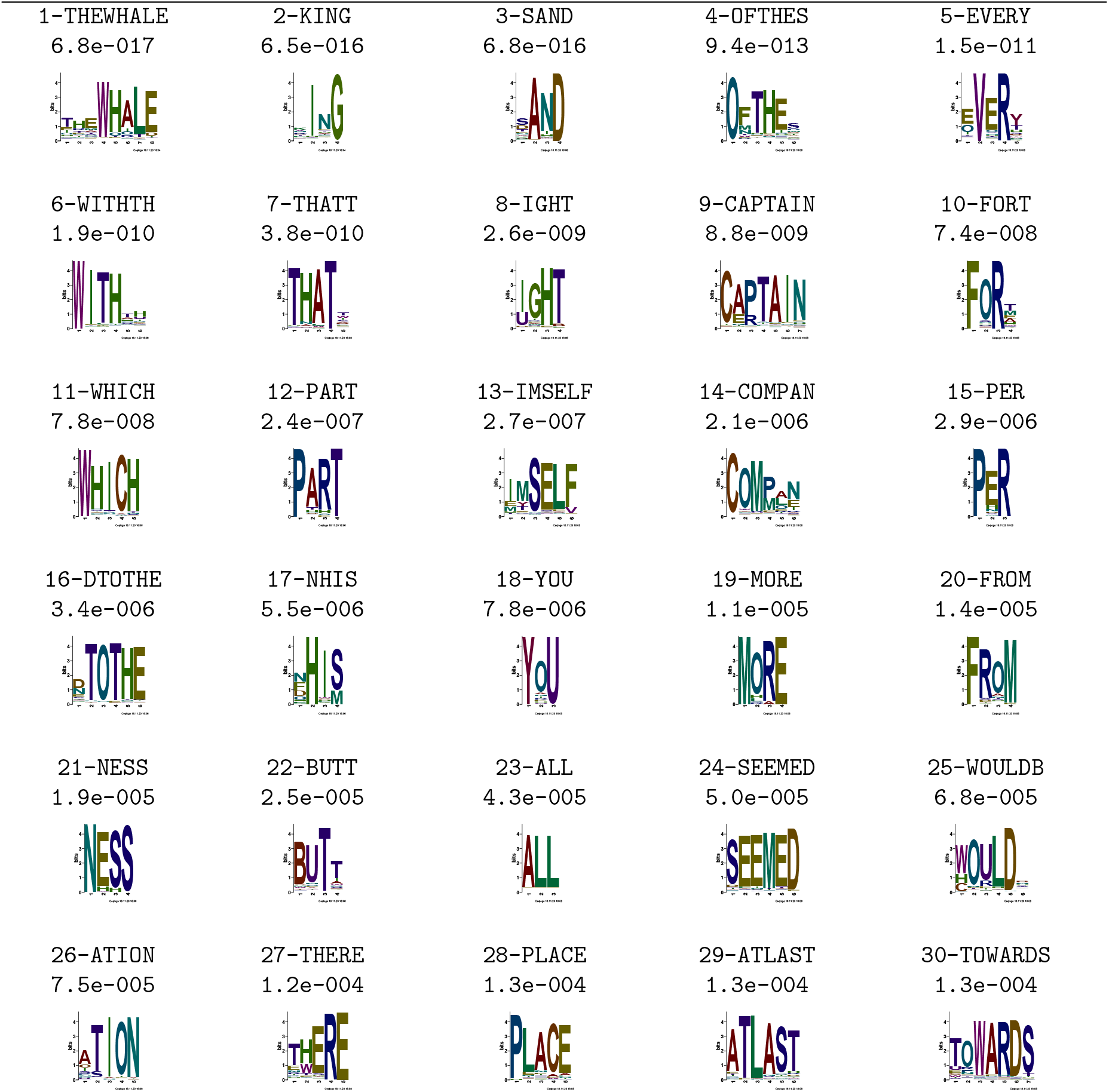
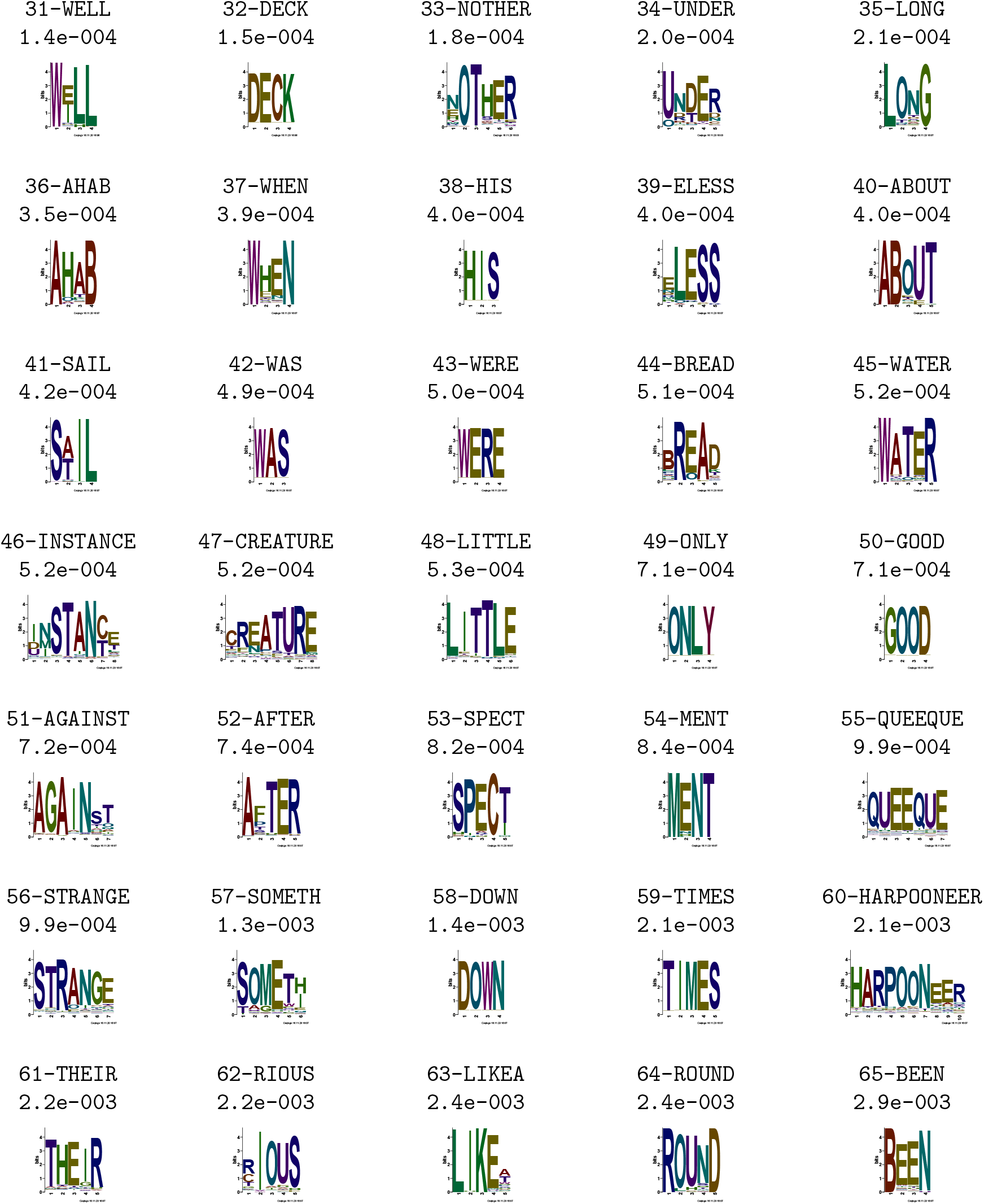
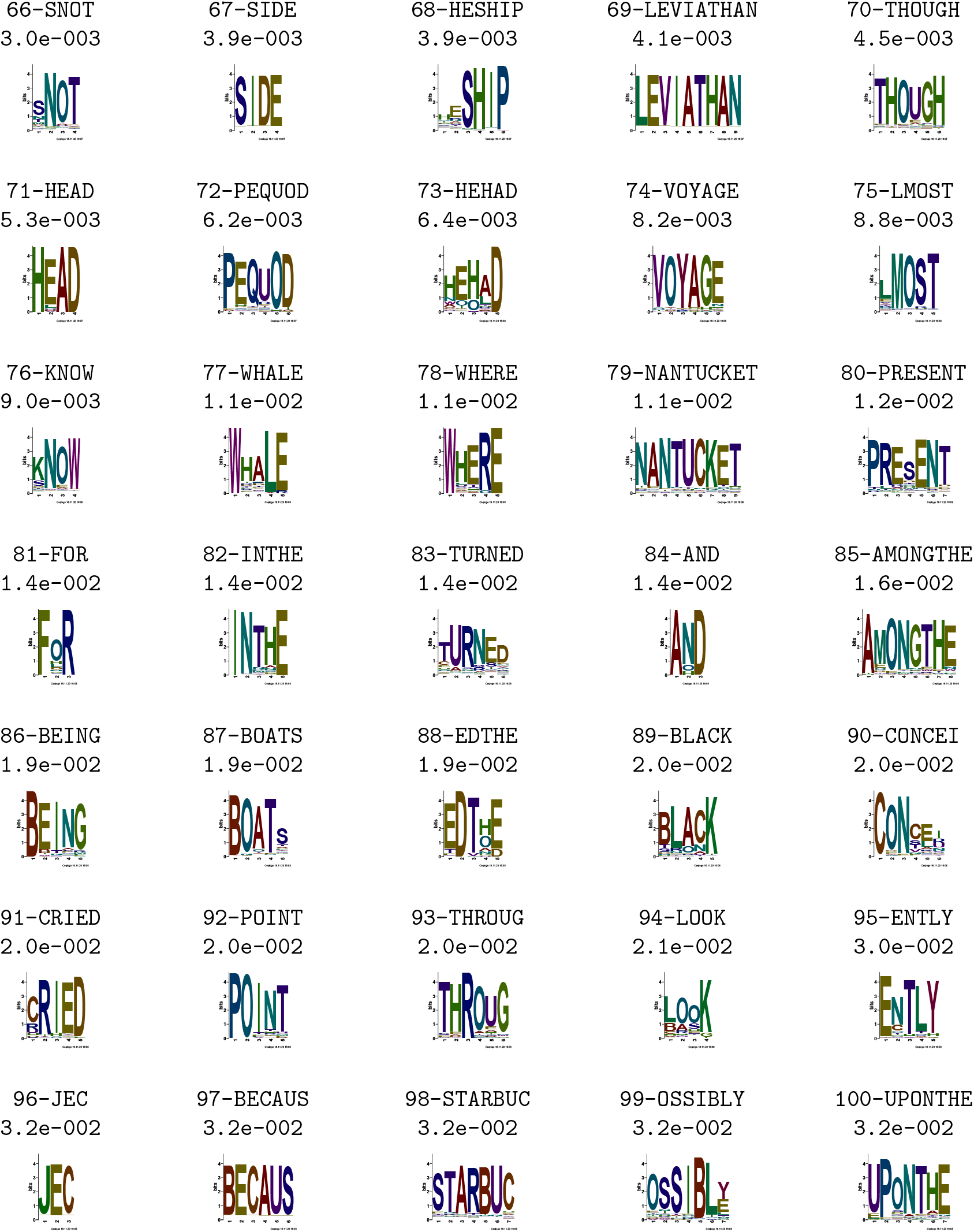
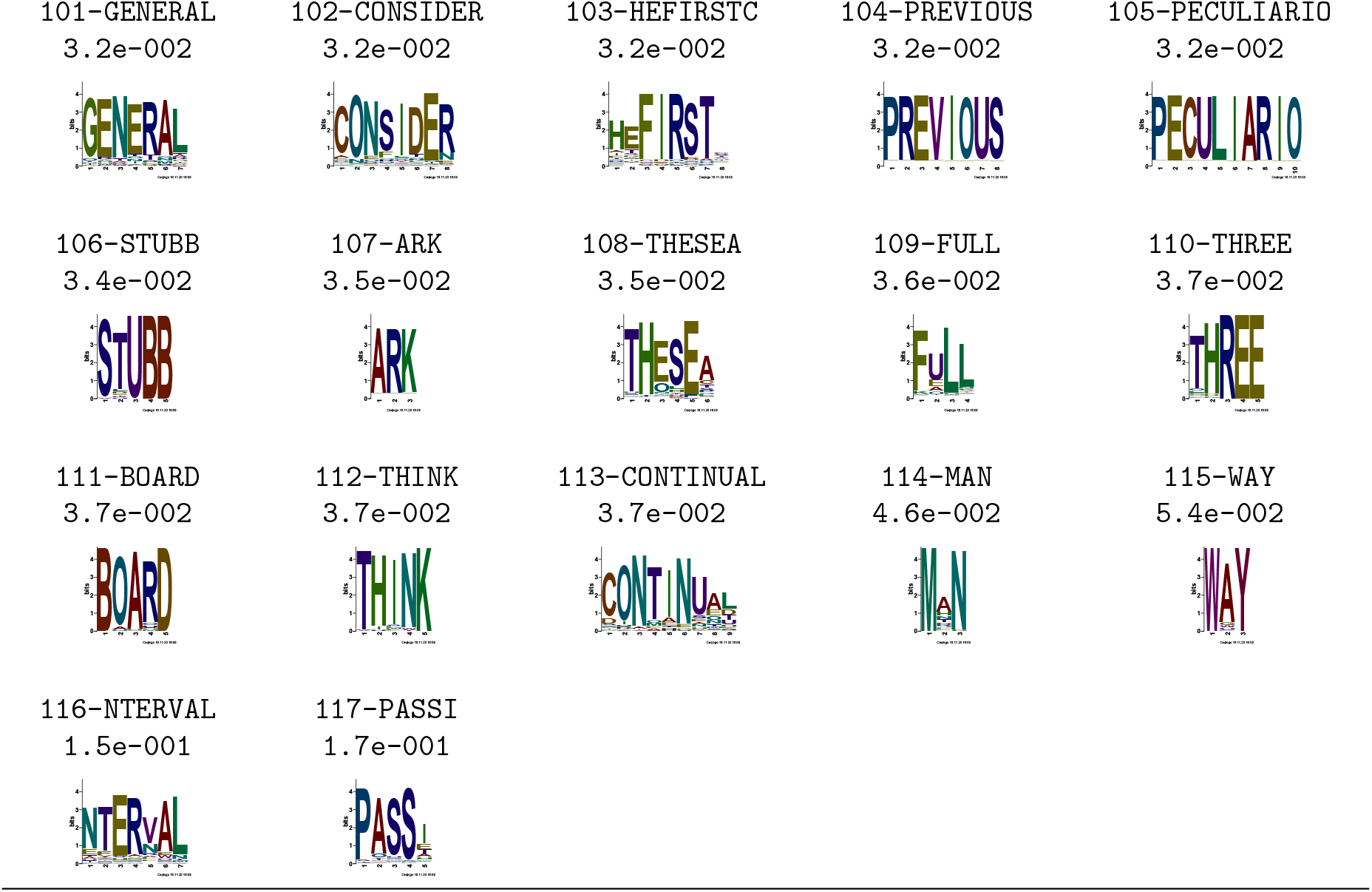
Motifs found by STREME in the corrupted text of the novel Moby Dick. The table shows names, *p*-values and sequence logos of the 117 motifs discovered by STREME in the text of Moby Dick, from which we removed all spacing, punctuation and capitalization, and to which we added 20% noise. We run STREME with the minimum motif width set to 3, the maximum width set to 12, and the size of *k*-mers to preserve set to 2.

The most significant motif STREME finds has the consensus sequence `THEWHALE`, which is certainly an extremely distinguishing phrase in this novel about whaling during the 19th century. The vast majority of the other motifs STREME discovers are either English words (e.g., `CAPTAIN, AHAB, LEVIATHAN`), English word suffixes (`ING, IOUS, ENTLY`), or short phrases (e.g., `OFTHE, INTHE, AMOUNGTHE`). For the most part, the motifs found by STREME begin and end precisely at correct word boundaries. Occasionally the first or last letter in the word or phrase is missing from the motif. An examination of these cases reveals that this is not a bug, but an artifact of the erasing of a prior motif site that slightly overlaps a site of the truncated motif (data not shown). STREME is thus effective at discovering informative motifs that capture interesting, distinguishing words and phrases in English text.

### 2.5 STREME accurately estimates motif statistical significance

Knowing the statistical significance of a discovered motif is a first step in deciding if it might by biologically significant. Several of the motif discovery algorithms studied here produce no estimates of motif significance (i.e., HOMER, Weeder, Peak-motifs), and the estimates produced by MEME can be extremely conservative, potentially causing biologically interesting motifs to be rejected by the user based on the estimate of statistical significance provided by MEME. DREME, provides accurate estimates of statistical significance, and we show here that STREME does, too.

For each motif STREME discovers, it reports an unbiased *p*-value that it estimates using a reserved portion of the sequences in its input. To verify the accuracy of these *p*-values, we run STREME on 1000 randomly generated datasets, each containing 10,000 sequences, allowing STREME to find exactly one motif in each dataset. Since all 1000 motifs are “discovered” in random sequences, their unbiased *p*-values should follow a uniform distribution. To check that they are uniformly distributed, we create a Q-Q plot [19], which plots the theoretical value of the *n*th largest *p*-value, *X* = 1/(*n* + 1), versus the *p*-value reported by STREME, *Y*.

The Q-Q plots in Fig. 9 show that the *p*-values reported by STREME are consistently accurate to within factor of 2. This is true for sequences over the DNA, RNA and protein alphabets, and regardless of whether the control sequences are the same length as the primary sequences (Fisher exact test) or not (Binomial test). The same holds true when we don’t provide a set of control sequences to STREME, causing it to create a control set by shuffling the primary sequences, and even when the input sequences have variable lengths from 50 to 150 characters (data not shown).

**Figure 9:**
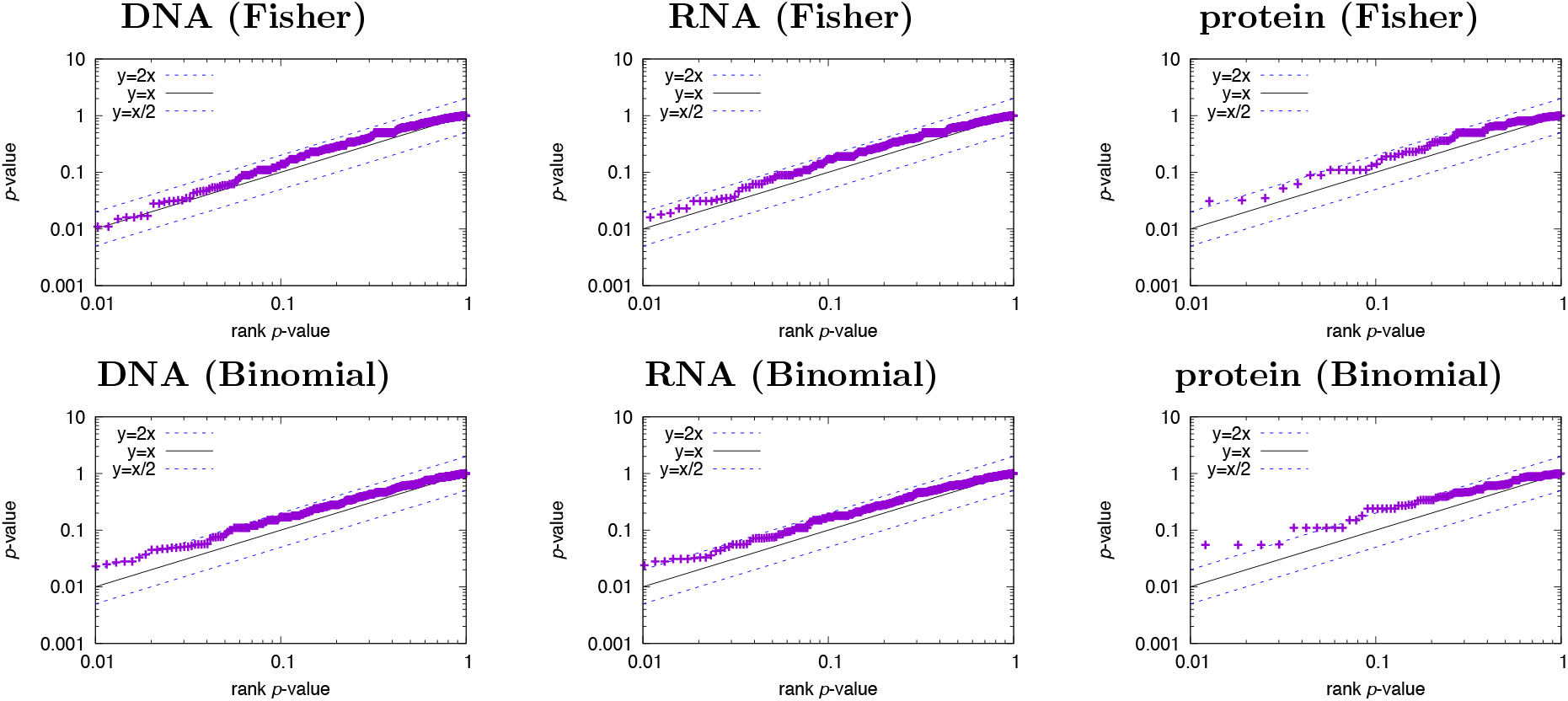
Q-Q accuracy plots of the *p*-values reported by STREME. Each panel shows the Q-Q plot for the *p*-values reported by STREME when run on 1000 datasets containing 10,000 random sequences over the alphabet given above the panel. Primary sequences are 100 characters long. Control sequences are 100 and 80 long in the first and second row of panels, respectively, causing STREME to use Fisher’s exact test (first row) or the Binomial test (second row). Ideally, the points should lie along the line *X* = *Y*.

We observe that the points in the Q-Q plots in Fig. 9 are consistently just above the line *X* = *Y*, showing that STREME’s *p*-values are slightly conservative. This means that if we accept STREME motifs with reported *p*-values of 0.05 or less, the Type I error (for each motif) will be no more than 5%, on average. Thus, the *p*-values reported by STREME can be used to judge whether an individual motif that it discovers is likely to be a false positive or not. Of course, when accepting multiple motifs, the user should apply multiple-testing correction such as a Bonferroni adjustment to the *p*-value threshold [14].

### 2.6 Speed of STREME with different alphabets and dataset sizes

We study the effect of the sequence alphabet and the size of the primary dataset using sets of artificially generated (random) sequences. We use datasets containing sequences of a fixed length, and adjust the number of sequences to create datasets containing from 100,000 to 20,000,000 characters (i.e., DNA or RNA bases or protein residues). We let STREME automatically create the control dataset, of equal size as the primary dataset. We allow STREME to report exactly five motifs, and let STREME pick the motif width between 5 and 30 positions. All runs are on a 3.2 GHz Intel Core i7 processor with 16GB of memory.

The results for DNA, RNA and protein sequences are shown in Fig. 10. Overall, the running time of STREME is roughly proportional to *n* log *n*, where *n* is the total primary dataset size in characters. For a given total primary sequence dataset size, running time tends to be maximal when the sequence length is approximately equal to the maximum allowed motif width, regardless of the alphabet size or the total dataset size.

**Figure 10:**
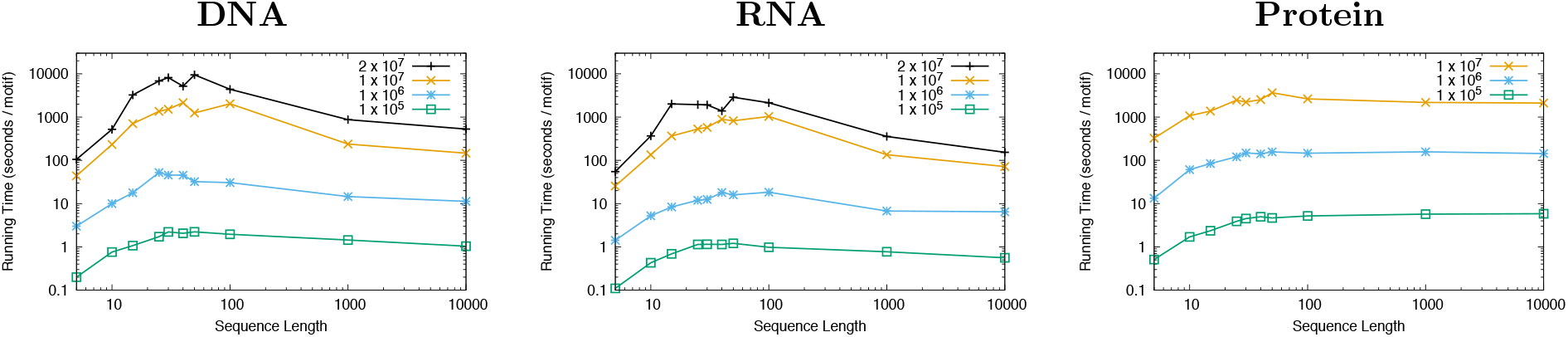
Running time of STREME on different types, lengths and numbers of sequences. Each point shows the running time in seconds per motif found (*Y*) when STREME run with a primary sequence dataset of a given *total size* (color), where all the sequences have the given length (*X*). The points for a given total size are connected with straight lines for ease of interpretation. The sequences were over the alphabet named above the panel, and were randomly generated uniformly using the MEME Suite `gendb` algorithm. STREME was run on a 3.2 GHz Intel Core i7 processor with 16GB of memory.

STREME is very fast, discovering motifs in 10,000 DNA sequences of length 100bp in about one minute per motif. With RNA sequences, STREME runs about twice as fast as with DNA since STREME treats RNA as single stranded, making the suffix tree half as large. Because the protein alphabet is larger, causing the suffix tree to have more branches, STREME runs about about 5 times more slowly with protein sequences than with RNA. In general, for single-stranded alphabets, the running time of STREME is roughly proportional to the size of the alphabet, and double-stranded alphabets take twice as long as a single-stranded alphabet of the same size.

## 3 Methods

### 3.1 STREME algorithm details

#### 3.1.1 Statistical tests of motif enrichment

When the primary and control sequences have identical length distributions, STREME uses the Fisher exact test to determine if a motif is enriched. Suppose that there *N_p_* and *N_c_* primary sequences and control sequences, respectively, and there are sites in *n_p_* and *n_c_* of the primary and control sequences, respectively. The Fisher exact test assumes a null model where a site is equally likely likely to be in any sequence. It computes the probability that *n_p_ or more* primary sequences would contain a site, and *n_c_ or fewer* control sequences would contain a site, given the values *N_p_, n_p_, N_c_* and *n_c_*.

When the primary and control sequence sets have different length distributions STREME uses the Binomial test (rather than Fisher’s exact test) to estimate the statistical significance of a discovered motif. Suppose that there are *N_p_* primary sequences with average length *L_p_*, and *N_c_* control sequences with average length *L_c_*. Then, on average, the number of possible positions a motif of width *w* could occupy is approximately *S_p_* = *N_p_*(*L_p_* – *w* – 1) and *S_c_* = *N_c_*(*L_c_* – *w* – 1), respectively. So STREME estimates the Bernoulli probability *P_b_* that a site chosen randomly in either of the two sets of sequences actually is in a primary sequence as *P_b_* = *S_p_/S_c_*. If *n_p_* primary sequences contain a match to a given motif (*n_p_* “successes”), the Binomial test considers the statistical significance of the motif to be the probability that *n_p_* or more primary sequences have matches to the motif. The test estimates the *p*-value of the motif as the sum of the binomial probabilities of *k* successes in *N_p_* trials, each with probability of success *P_b_*, where *k* goes from *n_p_* to *N_p_*,

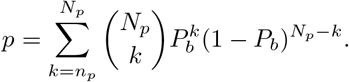

#### 3.1.2 The generalized suffix tree

STREME builds a generalized suffix tree from a single sequence that is the concatenation of all the input sequences (primary and control), with a unique “separator” character placed between sequences. Every suffix in the concatenated sequence corresponds to a leaf node, and vice-versa. The suffix for a leaf can be gotten by reading the labels on the branches along the path from the root to the leaf. STREME labels each leaf as to whether it corresponds to a suffix starting in primary or control sequence, and with the index of number of that sequence. The path from the root to any node in the tree corresponds to a word that occurs in the concatenated sequence. If the path does not contain the separator character, then the word actually occurs in one or more of the input sequences.

#### 3.1.3 Seed word evaluation

STREME adds to each node in the tree the counts of how many *sequences* (primary and secondary) contain the word corresponding to the node. It does this using a single pass of depth-first search, recording at each node the primary and control sequence index numbers present in the leaves below it. For “valid” nodes—those corresponding to actual words in the input sequences whose length lies in the desired motif width range—STREME computes a *p*-value by applying its enrichment test to the counts of primary and control sequences for the node. For each valid node, all its prefixes (of legal length) are called “initial seeds”.

STREME further evaluates the best 25 initial seeds of each legal width (best means lowest *p*-value). Evaluation of a seed begins with converting the word to a PWM using maximum likelihood estimation and a Dirichlet prior with a weight of 0.01. The Dirichlet prior is based on a 0-order Markov model of the control sequences. STREME then performs what we call “score-based matching” using the PWM and depth-first search of the suffix tree. STREME uses the PWM to compute the log-likelihood scores of the (legal-width) words corresponding to each node in the depth-first search. The search is pruned whenever the current node is too deep, or when the log-likelihood score of the word is below 0, indicating that the word is more likely under the background model than under the motif model. (Note that when STREME is using a higher-order Markov model of the control sequences, it uses the higher-order model along with the PWM to compute the correct log likelihood score of each word.) At the end of the search, STREME has recorded a list of nodes that correspond *approximate* matches to the PWM and to every prefix of the PWM. STREME uses these lists to determine the best matching site in each sequence. STREME also uses these lists to determine the counts of primary and control sequences that contain matches to the PWM, and computes the *p*-values of the counts for the PWM and all of its prefixes. Thus, in one pass of (pruned) depth-first search, STREME has scored 25 initial seeds and their prefixes as potential candidates for further refinement. These words are called “seed words”.

#### 3.1.4 Motif refinement

STREME further refines the four seed words of each legal width with the lowest *p*-values.

Refinement of a seed word begins by repeating the same steps as used above for evaluating initial seeds. STREME sorts the list of the best matching site in each sequence in order of decreasing log likelihood score, and finds the score threshold that yields the best enrichment *p*-value. STREME then estimates a new PWM (by maximum likelihood estimation) from just the sites above the threshold. These two steps— depth first search, reestimating the PWM—are iterated until the *p*-value fails to improve.

STREME selects the best final PWM from any of the refined seed words as the best motif, which it reports, and whose sites it then erases.

### 3.2 Evaluating motif discovery with TF ChIP-seq datasets

#### 3.2.1 Reference motifs

In order to evaluate the accuracy, sensitivity and speed of motif discovery algorithms on ChIP-seq datasets, we identify 40 TF ChIP-seq experiments in K562 cells for which there is a motif derived from high-throughput SELEX data for the same TF, or if not, for a member of the same transcription factor family. We download all 150 ENCODE AWG narrowPeak BED files of TF ChIP-seq experiments in K562 cells from the UCSC server at http://hgdownload.cse.ucsc.edu/goldenPath/hg19/encodeDCC/wgEncodeAwgTfbsUniform. Then, for each narrowPeak file, we create a FASTA file of 500bp sequences centered on each of the peaks. Next, using the CentriMo algorithm [8], we determine which of these ChIP-seq peak files shows central enrichment for high-throughput SELEX-derived motif [6]. For some TFs there are multiple SELEX motifs, and for others there is no motif for the TF itself, but there is a motif for a family member. For each of the 150 TF ChIP-seq experiments we assign the SELEX motif for the TF that is most highly enriched (if one exists) to the ChIP-seq peak file as the “reference” motif for that file. If there is no SELEX motif for the ChIP-ed TF, but there is one for a member of the TF’s family, we assign the most enriched family-member motif as the reference. These two rules allowed us to assign reference motifs to 40 of the 150 ENCODE TF ChIP-seq experiments. The names of the 40 ENCODE AWG BED files and the names of their reference motifs (which are available at http://meme-suite.org/meme-software/Databases/motifs/motif_databases.12.19.tgz in file `EUKARYOTE/Jolma2013.meme`) are given in Table 4.

**Table 4:**
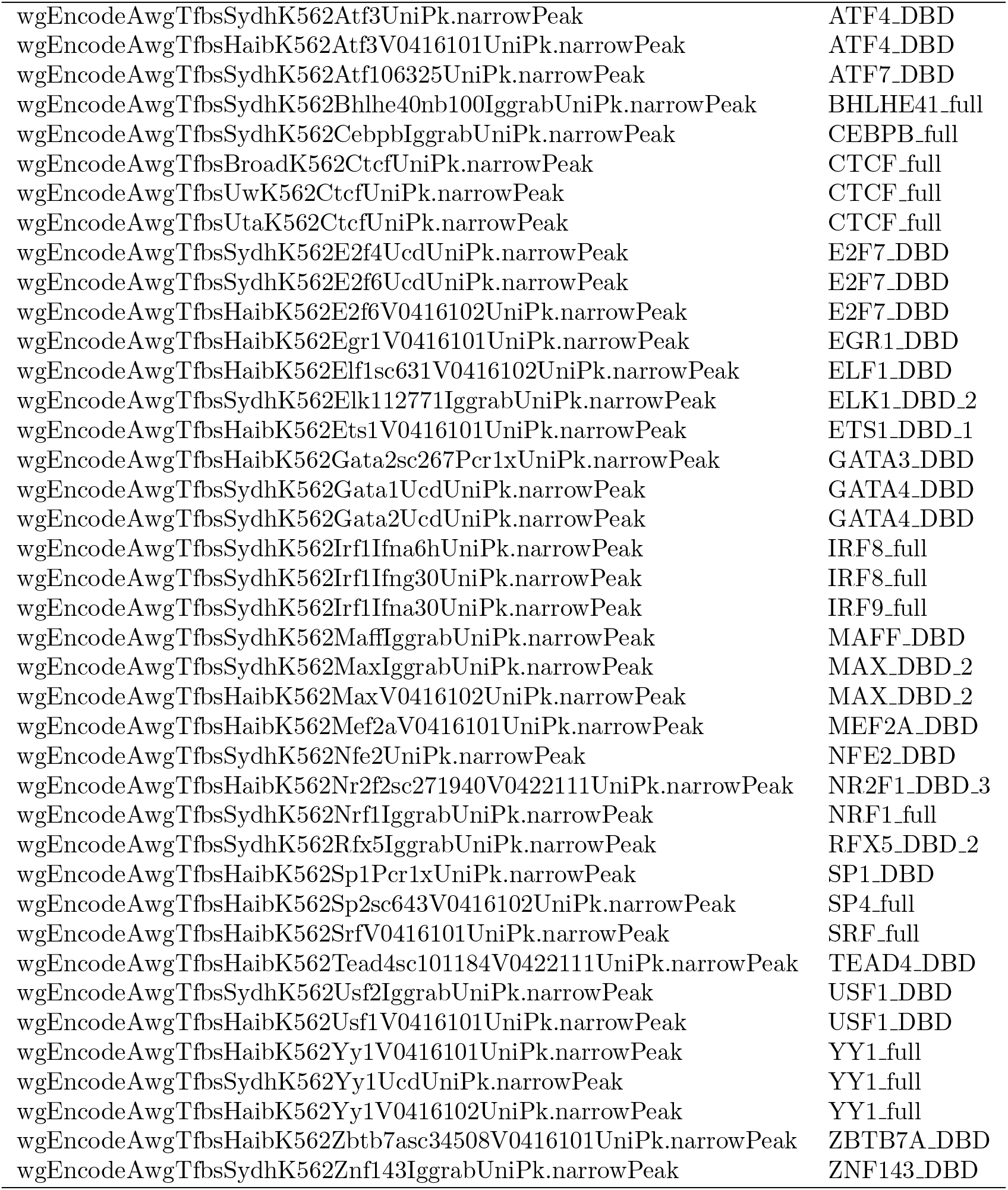
ENCODE K562 TF ChIP-seq files and SELEX [6] reference motifs. ChIP-seq files are available at http://hgdownload.cse.ucsc.edu/goldenPath/hg19/encodeDCC/wgEncodeAwgTfbsUniform with extension `gz.` Motifs are available at http://meme-suite.org/meme-software/Databases/motifs/motif_databases.12.19.tgz in file `EUKARYOTE/Jolma2013.meme`.

|  |  |
| --- | --- |
| wgEncodeAwgTfbsSydhK562Atf3UniPk.narrowPeak | ATF4_DBD |
| wgEncodeAwgTfbsHaibK562Atf3V0416101UniPk.narrowPeak | ATF4_DBD |
| wgEncodeAwgTfbsSydhK562Atf106325UniPk.narrowPeak | ATF7_DBD |
| wgEncodeAwgTfbsSydhK562Bhlhe40nb100IggrabUniPk.narrowPeak | BHLHE41_full |
| wgEncodeAwgTfbsSydhK562CebpbIggrabUniPk.narrowPeak | CEBPB_full |
| wgEncodeAwgTfbsBroadK562CtcfUniPk.narrowPeak | CTCF_full |
| wgEncodeAwgTfbsUwK562CtcfUniPk.narrowPeak | CTCF_full |
| wgEncodeAwgTfbsUtaK562CtcfUniPk.narrowPeak | CTCF_full |
| wgEncodeAwgTfbsSydhK562E2f4UcdUniPk.narrowPeak | E2F7_DBD |
| wgEncodeAwgTfbsSydhK562E2f6UcdUniPk.narrowPeak | E2F7_DBD |
| wgEncodeAwgTfbsHaibK562E2f6V0416102UniPk.narrowPeak | E2F7_DBD |
| wgEncodeAwgTfbsHaibK562Egr1V0416101UniPk.narrowPeak | EGR1_DBD |
| wgEncodeAwgTfbsHaibK562Elf1sc631V0416102UniPk.narrowPeak | ELF1_DBD |
| wgEncodeAwgTfbsSydhK562Elk112771IggrabUniPk.narrowPeak | ELK1_DBD_2 |
| wgEncodeAwgTfbsHaibK562Ets1V0416101UniPk.narrowPeak | ETS1_DBD_1 |
| wgEncodeAwgTfbsHaibK562Gata2sc267Pcr1xUniPk.narrowPeak | GATA3_DBD |
| wgEncodeAwgTfbsSydhK562Gata1UcdUniPk.narrowPeak | GATA4_DBD |
| wgEncodeAwgTfbsSydhK562Gata2UcdUniPk.narrowPeak | GATA4_DBD |
| wgEncodeAwgTfbsSydhK562Irf1Ifna6hUniPk.narrowPeak | IRF8_full |
| wgEncodeAwgTfbsSydhK562Irf1Ifng30UniPk.narrowPeak | IRF8_full |
| wgEncodeAwgTfbsSydhK562Irf1Ifna30UniPk.narrowPeak | IRF9_full |
| wgEncodeAwgTfbsSydhK562MaffIggrabUniPk.narrowPeak | MAFF_DBD |
| wgEncodeAwgTfbsSydhK562MaxIggrabUniPk.narrowPeak | MAX_DBD_2 |
| wgEncodeAwgTfbsHaibK562MaxV0416102UniPk.narrowPeak | MAX_DBD_2 |
| wgEncodeAwgTfbsHaibK562Mef2aV0416101UniPk.narrowPeak | MEF2A_DBD |
| wgEncodeAwgTfbsSydhK562Nfe2UniPk.narrowPeak | NFE2_DBD |
| wgEncodeAwgTfbsHaibK562Nr2f2sc271940V0422111UniPk.narrowPeak | NR2F1_DBD_3 |
| wgEncodeAwgTfbsSydhK562Nrf1IggrabUniPk.narrowPeak | NRF1_full |
| wgEncodeAwgTfbsSydhK562Rfx5IggrabUniPk.narrowPeak | RFX5_DBD_2 |
| wgEncodeAwgTfbsHaibK562Sp1Pcr1xUniPk.narrowPeak | SP1_DBD |
| wgEncodeAwgTfbsHaibK562Sp2sc643V0416102UniPk.narrowPeak | SP4_full |
| wgEncodeAwgTfbsHaibK562SrfV0416101UniPk.narrowPeak | SRF_full |
| wgEncodeAwgTfbsHaibK562Tead4sc101184V0422111UniPk.narrowPeak | TEAD4_DBD |
| wgEncodeAwgTfbsSydhK562Usf2IggrabUniPk.narrowPeak | USF1_DBD |
| wgEncodeAwgTfbsHaibK562Usf1V0416101UniPk.narrowPeak | USF1_DBD |
| wgEncodeAwgTfbsHaibK562Yy1V0416101UniPk.narrowPeak | YY1_full |
| wgEncodeAwgTfbsSydhK562Yy1UcdUniPk.narrowPeak | YY1_full |
| wgEncodeAwgTfbsHaibK562Yy1V0416102UniPk.narrowPeak | YY1_full |
| wgEncodeAwgTfbsHaibK562Zbtb7asc34508V0416101UniPk.narrowPeak | ZBTB7A_DBD |
| wgEncodeAwgTfbsSydhK562Znf143IggrabUniPk.narrowPeak | ZNF143_DBD |

**Table 5:** Non-redundant ENCODE K562 TF ChIP-seq files and SELEX [6] reference motifs. ChIP-seq files are available at http://hgdownload.cse.ucsc.edu/goldenPath/hg19/encodeDCC/wgEncodeAwgTfbsUniform with extension `gz.` Motifs are available at http://meme-suite.org/meme-software/Databases/motifs/motif_databases.12.19.tgz in file `EUKARYOTE/Jolma2013.meme`.

|  |  |
| --- | --- |
| wgEncodeAwgTfbsHaibK562Atf3V0416101UniPk.narrowPeak | ATF7_DBD |
| wgEncodeAwgTfbsSydhK562Bhlhe40nb100IggrabUniPk.narrowPeak | BHLHE41_full |
| wgEncodeAwgTfbsSydhK562CebpbIggrabUniPk.narrowPeak | CEBPB_full |
| wgEncodeAwgTfbsBroadK562CtcfUniPk.narrowPeak | CTCF_full |
| wgEncodeAwgTfbsHaibK562E2f6V0416102UniPk.narrowPeak | E2F7_DBD |
| wgEncodeAwgTfbsHaibK562Egr1V0416101UniPk.narrowPeak | EGR1_full |
| wgEncodeAwgTfbsHaibK562Elf1sc631V0416102UniPk.narrowPeak | ELF1_DBD |
| wgEncodeAwgTfbsHaibK562Gata2sc267Pcr1xUniPk.narrowPeak | GATA3_DBD |
| wgEncodeAwgTfbsSydhK562Irf1Ifna6hUniPk.narrowPeak | IRF8_full |
| wgEncodeAwgTfbsSydhK562MaffIggrabUniPk.narrowPeak | MAFF_DBD |
| wgEncodeAwgTfbsHaibK562Mef2aV0416101UniPk.narrowPeak | MEF2B_full |
| wgEncodeAwgTfbsSydhK562Nfe2UniPk.narrowPeak | NFE2_DBD |
| wgEncodeAwgTfbsHaibK562Nr2f2sc271940V0422111UniPk.narrowPeak | NR2F6_DBD_2 |
| wgEncodeAwgTfbsSydhK562Nrf1IggrabUniPk.narrowPeak | NRF1_full |
| wgEncodeAwgTfbsSydhK562Rfx5IggrabUniPk.narrowPeak | RFX5_DBD_2 |
| wgEncodeAwgTfbsHaibK562Sp1Pcr1xUniPk.narrowPeak | SP1_DBD |
| wgEncodeAwgTfbsHaibK562SrfV0416101UniPk.narrowPeak | SRF_full |
| wgEncodeAwgTfbsHaibK562Tead4sc101184V0422111UniPk.narrowPeak | TEAD4_DBD |
| wgEncodeAwgTfbsHaibK562Yy1V0416101UniPk.narrowPeak | YY1_full |
| wgEncodeAwgTfbsHaibK562Zbtb7asc34508V0416101UniPk.narrowPeak | ZBTB7B_full |
| wgEncodeAwgTfbsSydhK562Znf143IggrabUniPk.narrowPeak | ZNF143_DBD |

#### 3.2.2 Primary and control sequence datasets

For each of the 40 ENCODE TF ChIP-seq experiments, we create a single FASTA file containing the 100bp regions around the peaks by using the `fasta-fetch-centered tool` available in the MEME Suite software package to extract sequences from the *Homo sapiens* genome (hg19), which we download from the UCSC genome browser website (http://hgdownload.soe.ucsc.edu/goldenPath/hg19/bigZips/hg19.fa.gz). We use these files, which contain between 1,233 and 56,058 sequences, as the primary sequence datasets. For each such dataset, we create a control dataset by shuffling each sequence in the primary dataset using the `fasta-shuffle-letters` tool available in the MEME Suite software package. With this tool, the user can specify that the frequencies of words (*k*-mers) of any size *k* be preserved. The control dataset sequences are thus 100bp long, and have lower-order statistics matching those of the corresponding primary dataset.

To evaluate the thoroughness of motif discovery algorithms, we first identify a non-redundant subset of the 40 TF ChIP-seq sequence datasets described above, and we then create hybrid datasets containing sequences randomly selected from each of the 21 non-redundant TF ChIP-seq sequence datasets thus identified. To identify the non-redundant datasets, we use the `Tomtom` tool (available in the MEME Suite software package) to compare all 40 *reference motifs* to each other, and then greedily add motifs to the non-redundant reference set as long as their similarity to an already added motif is not too high (reject if Tomtom *p*-value ≤ 10^−4^). This results in a collection of 21 non-redundant reference motifs and their associated ChIP-seq datasets. We then create 20 hybrid primary sequence datasets by randomly selecting 100 sequences from each of the 21 non-redundant TF ChIP-seq dataset sequence files. (The sequences are all 100bp long.) Then, for each of the 20 hybrid primary datasets we create a matching control dataset by shuffling letters, as described above. The reason that we create the hybrid primary sequence datasets using only the non-redundant subset of ChIP-seq datasets is that we wish to avoid over-representing any motifs. Since many of the original 40 datasets contain ChIP-seq data for the same TF or for TFs in the same motif family, the reference motifs for some ChIP-seq datasets are very similar to each other. Our redundancy-filtering process eliminates this problem.

#### 3.2.3 Evaluating algorithm performance

To evaluate the accuracy, sensitivity and thoroughness of the motif discovery algorithms, we compare the motifs they discover to all 843 motifs in the Jolma compendium of SELEX-derived motifs [6] using the Tomtom tool. Our figure of merit is the best (minimum) Tomtom *p*-value of the similarity between the SELEX reference motif for the ChIP-seq dataset and the motifs reported by the algorithm. For a range of similarity (motif accuracy) thresholds, we count the number of times each algorithm finds a motif at least that similar to the SELEX reference motif for the ChIP-seq dataset. For ease of exposition, we convert the Tomtom *p*-value to our “motif accuracy score”, which is minus the base-10 logarithm of the *p*-value.

### 3.3 Evaluating motif discovery with CLIP-seq datasets

We follow a similar procedure to the above in identifying and preparing a group of CLIP-seq datasets and reference motifs. The datasets are derived from ENCODE enhanced CLIP-seq (eCLIP) databases from the Yeo lab [17], and the reference motifs come from the RNAcompete database [11]. We manually determine the set of 20 RNA-binding proteins with ENCODE eCLIP datasets from K562 cells that also have (at least one) motif in the RNAcompete compendium for *Homo sapiens*. We then download the eCLIP narrowPeak BED files that combine two eCLIP replicates from the ENCODE website (https://www.encodeproject.org/search) for those 20 experiments. From each of the 20 BED files we create a BED file of non-overlapping regions using a custom script. From each non-overlapping BED file we create a FASTA file of sequences using the `fasta-fetch-centered` tool from MEME Suite specifying its `-r` option in order to fetch the exact regions specified in the BED file from the *Homo sapiens* hg19 genome. We use the 20 FASTA files thus created, which contain between 166 and 7,520 sequences with average length 70, as the primary sequence datasets in our evaluation of motif discovery algorithms. The ENCODE accession numbers and reference motifs for the eCLIP datasets are given in Table 6.

**Table 6:**
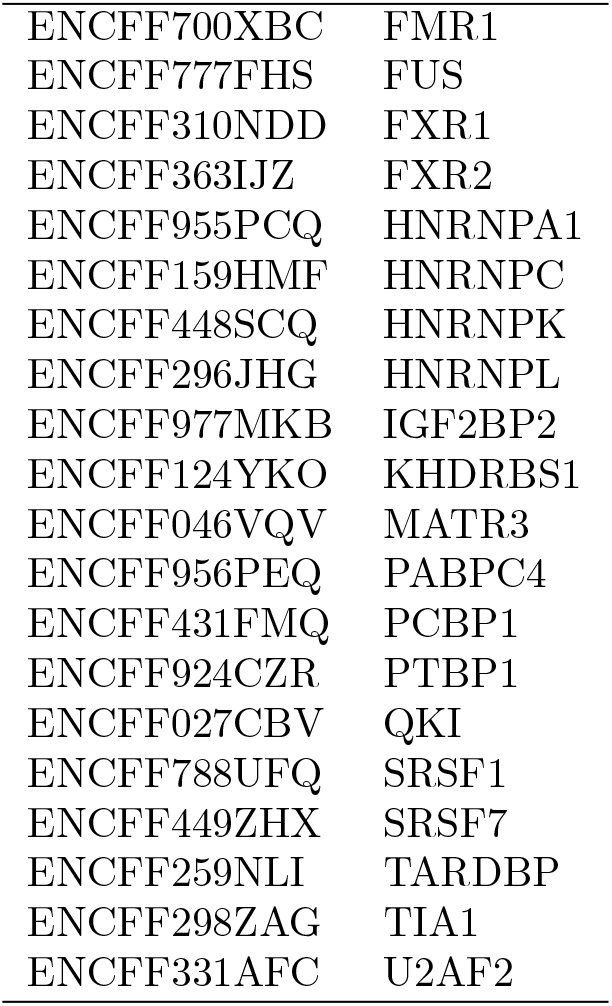
ENCODE K562 eCLIP files and RNAcompete [11] reference motifs. eCLIP narrowPeak files are available at https://www.encodeproject.org/search. Motifs are available at http://meme-suite.org/meme-software/Databases/motifs/motif_databases.12.19.tgz in file `RNA/Ray2013xrbp_Homo_sapiens.meme`.

|  |  |
| --- | --- |
| ENCFF700XBC | FMR1 |
| ENCFF777FHS | FUS |
| ENCFF310NDD | FXR1 |
| ENCFF363IJZ | FXR2 |
| ENCFF955PCQ | HNRNPA1 |
| ENCFF159HMF | HNRNPC |
| ENCFF448SCQ | HNRNPK |
| ENCFF296JHG | HNRNPL |
| ENCFF977MKB | IGF2BP2 |
| ENCFF124YKO | KHDRBS1 |
| ENCFF046VQV | MATR3 |
| ENCFF956PEQ | PABPC4 |
| ENCFF431FMQ | PCBP1 |
| ENCFF924CZR | PTBP1 |
| ENCFF027CBV | QKI |
| ENCFF788UFQ | SRSF1 |
| ENCFF449ZHX | SRSF7 |
| ENCFF259NLI | TARDBP |
| ENCFF298ZAG | TIA1 |
| ENCFF331AFC | U2AF2 |

We find that using a shuffled version of the primary sequence dataset as the control dataset does not work as well with this eCLIP data and these reference motifs as using a more random control dataset. All the motif discovery algorithms we test here perform better when we construct a single control dataset by consisting of 10,000 sequences randomly chosen from the combined sequences in the 20 primary sequence datasets. This is the control dataset we use here in all eCLIP-based analyses.

We evaluate algorithm performance on the eCLIP datasets analogously to our approach with the ChIP-seq datasets (see previous section).

### 3.4 Evaluating STREME with a custom alphabet

To evaluate STREME using a custom alphabet, we create a “corrupted” version of the novel “Moby Dick”. We download the complete text of the novel from the Gutenberg Project (https://gutenberg.org). To make motif discovery more difficult, we remove all punctuation, spaces and capitalization from the text. We then convert the text to FASTA format, placing each line in the original text file in a single FASTA sequence. To make the problem more similar to a biological one, we then corrupt the text by randomly changing 20% of the letters in each FASTA sequence to a different, randomly-chosen letter. We then run STREME on the resulting FASTA file, along with the alphabet definition file that defines the 26 letter alphabet of the sequences.

### 3.5 Creating random sequence datasets

To evaluate the accuracy of STREME *p*-values, and to evaluate speed of STREME on different alphabets and dataset sizes, we use randomly generated sequences. We generate random DNA and protein sequence datasets using the `gendb` tool from the MEME Suite package. This tool allows us to specify the number as well as the minimum and maximum length of generated sequences. We generate RNA sequences by replacing `T` with `U` in DNA sequences generated by `gendb`.

## 4 Discussion

Our results show that STREME is a versatile new algorithm for *ab initio* discovery of sequence motifs in very large sequence datasets. With transcription factor ChIP-seq data, our study indicates that STREME usually finds more accurate motifs than the other algorithms with which we compare it (HOMER, MEME, Weeder, DREME and Peakmotifs). STREME also appears to be better at discovering the motifs of other transcription factors that bind cooperatively with, or in proximity to, the ChIP-ed TF. STREME is generally faster than the other algorithms we study here, especially when finding motifs wider than 12 positions.

Our study also suggest that STREME performs as well as other motif discovery algorithms when applied to RNA from CLIP-seq experiments. In addition to DNA and RNA sequences, STREME can be applied to motif discovery in protein sequences, although we do not study that application here. Additionally, STREME allows the user to define their own sequence alphabet, and will discover motifs present in sequences over that alphabet. We illustrate this capability by using STREME to “discover” the key words and phrases in a highly corrupted version of an English text.

With datasets containing fewer than 100 sequences, our results show that STREME is not quite as accurate as MEME, but is more accurate than the other motif discovery algorithms in our study. We intend to incorporate STREME into the MEMEChIP algorithm, which performs a comprehensive analysis of sequences from ChIP-seq (and similar) experiments. Currently, MEME-ChIP combines the motifs discovered by MEME and DREME into a single, non-redundant analysis. STREME will replace DREME in that role. As our results show, STREME outperforms DREME in all the respects we study here—accuracy, sensitivity, thoroughness and speed—so we expect this change to enhance those aspects of MEME-ChIP as well.

**Figure 11:**
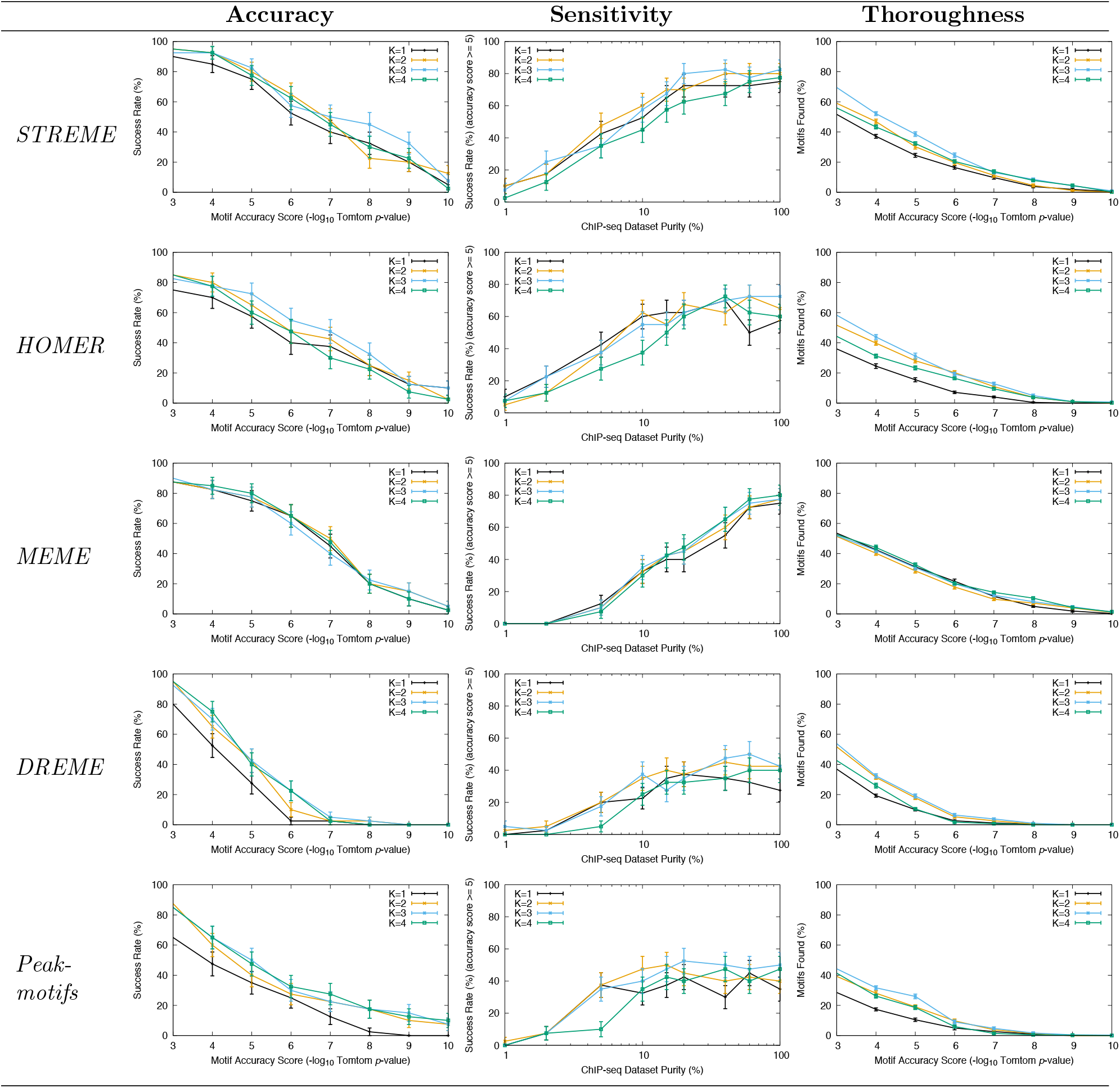
Choice of shuffling *k*-mer size (*k*) on motif discovery in ChIP-seq datasets. The panels show the accuracy, sensitivity and thoroughness of different motif discovery algorithms as a function of the size of words (*k*) whose frequencies are preserved when shuffling the primary sequences to create the control sequences. STREME, MEME and Peak-motifs also use *k* – 1 as the order of the Markov model they create internally from the input sequences. (Weeder does not use a control set.) The contents of the plots in the three columns are described in the captions to Figures 1, 2 and 3, respectively. The sensitivity plots use a motif accuracy score threshold of 5 (Tomtom *p*-value ≤ 10^−5^).

## 5 Acknowledgments

I thank Charles E. Grant for proofreading the manuscript.

## References

[1] T. L. Bailey. DREME: motif discovery in transcription factor ChIP-seq data. Bioinformatics, 27(12):1653–1659, Jun 2011.

[2] T. L. Bailey and C. Elkan. The value of prior knowledge in discovering motifs with MEME. Proceedings of the Third International Conference on Intelligent Systems for Molecular Biology, Cambridge, United Kingdom, July 16-19, 1995, 3:21–29, 1995.

[3] R. A. Fisher. On the interpretation of χ^2^ from contingency tables, and the calculation of p. Journal of the Royal Statistical Society, 85(1):87–94, 1922.

[4] S. Gupta, J. Stamatoyannopoulos, T. Bailey, and W. S. Noble. Quantifying similarity between motifs. Genome Biol, 8(2):R24, Feb 2007.

[5] S. Heinz, C. Benner, N. Spann, E. Bertolino, Y. C. Lin, P. Laslo, J. X. Cheng, C. Murre, H. Singh, and C. K. Glass. Simple combinations of lineage-determining transcription factors prime cis-regulatory elements required for macrophage and B cell identities. Molecular cell, 38:576–589, May 2010.

[6] A. Jolma, J. Yan, T. Whitington, J. Toivonen, K. R. Nitta, P. Rastas, E. Morgunova, M. Enge, M. Taipale, G. Wei, K. Palin, J. M. Vaquerizas, R. Vincentelli, N. M. Luscombe, T. R. Hughes, P. Lemaire, E. Ukkonen, T. Kivioja, and J. Taipale. DNA-binding specificities of human transcription factors. Cell, 152(1-2):327–339, Jan 2013.

[7] S. Kurtz, A. Phillippy, A. L. Delcher, M. Smoot, M. Shumway, C. Antonescu, and S. L. Salzberg. Versatile and open software for comparing large genomes. Genome biology, 5:R12, 2004.

[8] T. Lesluyes, J. Johnson, P. Machanick, and T. L. Bailey. Differential motif enrichment analysis of paired ChIP-seq experiments. BMC Genomics, 15:752, 2014.

[9] N. Nagarajan, N. Jones, and U. Keich. Computing the P-value of the information content from an alignment of multiple sequences. Bioinformatics, 21 Suppl 1:i311–i318, Jun 2005.

[10] G. Pavesi, P. Mereghetti, G. Mauri, and G. Pesole. Weeder Web: discovery of transcription factor binding sites in a set of sequences from co-regulated genes. Nucl Acids Res, 32(Web Server issue):W199–W203, Jul 2004.

[11] D. Ray, H. Kazan, K. B. Cook, M. T. Weirauch, H. S. Najafabadi, X. Li, S. Gueroussov, M. Albu, H. Zheng, A. Yang, H. Na, M. Irimia, L. H. Matzat, R. K. Dale, S. A. Smith, C. A. Yarosh, S. M. Kelly, B. Nabet, D. Mecenas, W. Li, R. S. Laishram, M. Qiao, H. D. Lipshitz, F. Piano, A. H. Corbett, R. P. Carstens, B. J. Frey, R. A. Anderson, K. W. Lynch, L. O. F. Penalva, E. P. Lei, A. G. Fraser, B. J. Blencowe, Q. D. Morris, and T. R. Hughes. A compendium of RNA-binding motifs for decoding gene regulation. Nature, 499(7457):172–177, Jul 2013.

[12] J. E. Reid and L. Wernisch. STEME: efficient EM to find motifs in large data sets. Nucl Acids Res, 39(18):e126, Oct 2011.

[13] T. D. Schneider and R. M. Stephens. Sequence logos: a new way to display consensus sequences. Nucl Acids Res, 18(20):6097–6100, Oct 1990.

[14] R. J. Simes. An improved Bonferroni procedure for multiple tests of significance. Biometrika, 73:751–754, 1986.

[15] G. D. Stormo. DNA binding sites: representation and discovery. Bioinformatics, 16(1):16–23, Jan 2000.

[16] M. Thomas-Chollier, M. Defrance, A. Medina-Rivera, O. Sand, C. Herrmann, D. Thieffry, and J. van Helden. RSAT 2011: regulatory sequence analysis tools. Nucl Acids Res, 39(Web Server issue):W86–W91, Jul 2011.

[17] E. L. Van Nostrand, G. A. Pratt, A. A. Shishkin, C. Gelboin-Burkhart, M. Y. Fang, B. Sundararaman, S. M. Blue, T. B. Nguyen, C. Surka, K. Elkins, R. Stanton, F. Rigo, M. Guttman, and G. W. Yeo. Robust transcriptome-wide discovery of RNA-binding protein binding sites with enhanced CLIP (eCLIP). Nature methods, 13:508–514, June 2016.

[18] P. Weiner. Linear pattern matching algorithms. In Ifth Annual Symposium on Switching and Automata Theory, pages 1–11. IEEE, 1973.

[19] M. B. Wilk and R. Gnanadesikan. Probability plotting methods for the analysis of data. Biometrika, 55(1):1–17, 03 1968.

